# Comparative analysis of microglia-targeted AAVs reveals capsid choice drives efficiency *in vitro* but has limited impact *in vivo*

**DOI:** 10.64898/2026.08.05.739854

**Authors:** Saša Jereb, Lia R. D’Alessandro, Nader Morshed, Margery Chen, Rafaela Sartore, Zengpeng Han, Jordan E. McKinney, Pamela P. Brauer, John W. Harvey, Matthew Demers, Dara Cuffe, Raphael Rakosi-Schmidt, Luira Leite, Youtong Huang, Qingxia Zheng, Chin-Yen Lin, Ken Y. Chan, Bryan J. Song, Michael R. Farzan, Paola Arlotta, Morgan Sheng, Mariko L. Bennett, Matthew B. Johnson, Beth Stevens, Benjamin E. Deverman

**Author notes:** These authors contributed equally (Beth Stevens, Benjamin E. Deverman).

## Abstract

Microglia play key roles in brain development, homeostasis, and neurodegeneration. Although multiple strategies for viral gene delivery to microglia have been reported, they have not been directly compared. Here, we developed microglia-targeting AAV capsids and benchmarked them against existing approaches. The novel capsids exhibit improved transduction efficiency in cultured mouse and human microglia, as well as neurons and astrocytes. However, when we compared microglial transduction efficiency of the novel capsids with published engineered and naturally occurring capsids after intracranial injection, all capsids achieved efficient and specific transduction when paired with a genome incorporating *IBA1* promoter and miR-124 target sites. In contrast, CAG promoter did not support efficient microglial transduction. Moreover, blood-brain barrier-crossing capsids carrying *IBA1* promoter and miR-124 target sites efficiently transduced microglia at high doses but exhibited off-target expression. Together, our work provides improved capsids for *in vitro* manipulation of microglia and establishes viral genome design, not capsid identity, as the principal determinant of efficient *in vivo* microglial targeting.

## Introduction

Microglia play essential roles in brain health and disease. Genome-wide association and exome-sequencing studies have shown that multiple Alzheimer’s disease (AD) risk variants are located in or near microglia-expressed genes (Kunkle et al., 2019). Microglia also adopt distinct molecular states in response to AD and other brain pathologies (Keren-Shaul et al., 2017; Krasemann et al., 2017), further supporting their central role in disease pathogenesis. Beyond disease, microglia are critical responders to brain injury and infection, as well as important players in brain development (Favuzzi et al., 2021; Schafer et al., 2012). Single-cell studies have revealed their extensive heterogeneity across development and ageing, including axon-tract associated microglia (Hammond et al., 2019; Li et al., 2019) and white-matter associated microglia (Safaiyan et al., 2021). Together, these findings underscore the need for scalable tools to interrogate gene function across microglial states and contexts.

Adeno-associated viruses (AAVs) enable rapid gene manipulation across species and offer a promising platform for high-throughput microglial perturbation *in vitro* and *in vivo*. Microglia-targeting AAVs could also support development of gene therapies for disorders caused by mutations in microglia-expressed genes, e.g. *CSF1R* (Rademakers et al., 2011), *SAMHD1* (Rice et al., 2009) and *TREM2/TYROBP* (Paloneva et al., 2002). While microglia were believed to be refractory to viral transduction (Maes et al., 2019), multiple recent papers reported improved microglia targeting *in vivo* in mice either through capsid (Lin et al., 2022; Santoscoy et al., 2024) or genome engineering (Aoki et al., 2025; Okada et al., 2022; Serrano et al., 2024). However, these approaches have not been systematically compared, and their reproducibility remains uncertain (Lin et al. 2022 study has been challenged by Cao et al., 2025), limiting a clear understanding of variables that affect efficient and specific targeting and preventing widespread adoption of these tools by the field.

Here, we sought to determine the relative contributions of capsid engineering and viral genome design to efficient and specific microglial transduction by comparing newly engineered and previously reported AAV strategies. We first tested whether capsid engineering could enhance microglial transduction, motivated by our observation that AAV9 genomes do not accumulate in microglial cells following intracranial injection, suggesting a bottleneck at the level of cellular entry or intracellular trafficking. We screened AAV9 capsid libraries for their ability to transduce microglia *in vitro* and *in vivo*. *In vitro*, we identified variants with enhanced transduction of mouse primary microglia and human embryonic stem cell (ESC)-derived microglia (iMGLs) (Abud et al., 2017) with significantly higher efficiency than AAV9. However, systematic *in vivo* comparison of our engineered capsids with previously published microglia-targeting engineered and naturally occurring capsids revealed that capsid identity is not the key determinant of microglial transduction efficiency following intracranial injection. Rather, it is the transgene regulatory elements that determine transduction efficiency of microglia. Finally, we demonstrated that intravenous administration of blood-brain barrier (BBB)-crossing capsids can lead to microglial transduction at high doses but resulted in substantial off-target expression. Together, our findings establish that viral genome design, rather than capsid identity, is the principal determinant of efficient and specific microglial transduction *in vivo* while providing improved capsids for *in vitro* genetic manipulation.

## Results

### AAV genomes preferentially accumulate in non-microglia nuclei

AAV particles, which carry a single-stranded DNA genome, are endocytosed and trafficked through endolysosomal vesicles and the trans-Golgi network, then escape into the cytosol and enter the nucleus (Nonnenmacher & Weber, 2011; Pillay et al., 2016). There, the capsid uncoats and the second DNA strand is synthesized before transcription can occur (Berry & Asokan, 2016). To understand the factors limiting microglia transduction by AAVs, we first asked whether AAV DNA can be detected in microglia *in vivo*. This would help us answer whether AAV entry and trafficking to the nucleus is a significant bottleneck. We intravenously injected adult wild-type (WT) mice with AAV- PHP.B, a BBB-crossing capsid (Deverman et al., 2016), carrying a CAG-GFP-NLS genome. After 3 weeks, we performed FACS to sort microglia and astrocytes from the injected mice, followed by PCR amplification of AAV genomes (**Supplementary Fig. 1a**). We found significantly fewer AAV genomes in microglia compared to astrocytes (**Supplementary Fig. 1b**). However, compared to the negative control (microglia from mice not injected with AAV-PHP.B), we detected more AAV genomes in microglia from injected mice, which left open the question of whether the AAV genomes we detected were intracellular or originated from debris from transduced cells that co-purified with microglia.

To investigate this question in more detail, we used *in situ* hybridization to detect AAV DNA in microglia after intracranial injection of AAV9. We injected AAV9 carrying a CAG-GFP-NLS genome into the striatum of adult mice. We sacrificed and perfused mice at 6 hours, 9 hours, and 18 hours after injection (**Supplementary Fig. 1c**). We then performed RNAscope to detect AAV genomes (Zhao et al., 2020), followed by immunofluorescence for Iba1, a microglial marker. To specifically detect AAV DNA, and not AAV RNA, we used a probe set designed to hybridize to the CMV enhancer present in the AAV genome. At 6 hours after injection, AAV genomes were predominantly extracellular. By 9 and 18 hours the AAV genomes accumulated at high density within non-microglial (Iba1 negative) nuclei, but were virtually undetectable within Iba1 positive microglia. This finding suggests that few if any AAV genomes accumulate in microglia (**Supplementary Fig. 1d**).

We also tested the engineered microglia-targeting capsids AAV-MG1.1 and AAV- MG1.2 (Lin et al., 2022). These capsids were reported to mediate highly efficient and specific microglia transduction when injected intracranially into Cx3cr1-Cre-ER mice that express Cre in microglia (Parkhurst et al., 2013) in combination with an AAV genome comprising the SFFV promoter and a Cre-dependent system for reporter gene expression (DIO/FLEx system). A recent study has failed to replicate the high microglia transduction efficiency of the MG1.2 capsid (Cao et al., 2025). However, the experimental conditions in this study were different from the ones reported in the original publication, as the authors used the CAG promoter and injected AAV-MG1.2 into WT mice (Cao et al., 2025). In our experiment, we matched the conditions reported in the Lin et al. paper as closely as possible, but we were still not able to replicate the finding (**Supplementary Fig. 2a**). We stained the injected brains with an *in-situ* hybridization probe set binding to WPRE in the AAV DNA to confirm that our intracranial injections were successful as well as to check whether the AAV genomes packaged in the engineered capsids are internalized by microglia. Similar to the initial time course data using AAV9 (**Supplementary Fig. 1b**), AAV genomes delivered by the MG1.1 and MG1.2 capsids accumulated in non-microglial cells, but not microglia (**Supplementary Fig. 2b**).

These data suggested that limited viral entry or intracellular trafficking represented the major early barrier to microglial transduction, motivating us to test whether capsid engineering could overcome this bottleneck.

### *In vitro* screening identifies AAV capsids that transduce cultured mouse microglia with high efficiency

We performed an *in vivo* capsid selection using Cre recombination-based AAV targeted evolution (CREATE), which selects for AAV particles that are internalized and have their genome converted to double-stranded DNA within Cre expressing cells using Cre recombination-dependent PCR (Deverman et al., 2016). For this experiment, we used Cx3cr1-Cre-ER mice to provide selective pressure for capsids that transduce Cre+ microglia (Parkhurst et al., 2013) and WT mice as controls. We injected the mice intracranially in the cortex and striatum with an AAV capsid library containing random 7- mer amino acid insertions in loop VIII of the AAV9 capsid protein. However, when we analyzed the sequences recovered from these mice, the correlation between the mean reads per million (RPM) of the most abundant capsid sequences from WT control and Cre+ animals was high (R=0.83 for the 100 most abundant sequences from Cre+ mice), suggesting that either a wide variety of sequences were able to enter microglia *in vivo* or that the selective pressure was too stringent and the only recovered sequences were rare AAV genomes with lox sites that had spontaneously recombined in the initial virus library.

We therefore shifted our strategy to an *in vitro* capsid selection to identify capsids that are highly efficient at binding and transducing microglia *in vitro* but could potentially also transduce microglia *in vivo*. Unlike the *in vivo* selection, this approach enabled us to expose the capsid library at a low multiplicity of infection (10K vg/cell) to a large number of microglia (10 million per replicate), resulting in high selective pressure and high signal to noise ratio. For the selection, we used cultured mouse microglia (isolated from mixed culture with astrocytes) and bone marrow macrophages (BMMs, an additional macrophage cell type to compare with microglia), as well as cultured neurons, astrocytes, and HEK cells to identify capsids that are enriched in microglia/macrophages compared to other cell types (**Fig. 1a**). Importantly, we FACS-purified microglia from mixed cultures to remove astrocytes that could lead to false positives. To provide two different levels of selective pressure, we performed both binding and transduction assays for each cell type. Given that microglia and BMMs are difficult to transduce *in vitro* with WT AAV9, binding assays provide a less stringent selection. For the binding assay, we exposed cells to the AAV library for 2 hours at 4°C, washed and sequenced the AAV capsid DNA to select for capsids with increased binding to each cell type. For the transduction assays, we exposed the cells to AAV, harvested RNA 5 days later, and sequenced AAV cDNA to identify capsids that successfully transduced the cells. In line with our expectation, we recovered more sequences from the binding assays compared to the transduction assays (from microglia samples, we recovered a total of 294,037 vs. 106,803 sequences represented by at least one sequencing read, respectively). Interestingly, 80 of the top 100 capsid variants that bound microglia also showed high binding to bone marrow macrophages (defined as average RPM >150), whereas only 58 and 40 variants showed high binding to neurons and astrocytes, respectively. Similarly, when we considered top sequences that were specifically bound to microglia (defined as those where the ratio of mean RPM from microglia over mean RPM from non-macrophage samples is higher than 4), we found that these sequences were also highly abundant among sequences bound to BMMs (**Fig. 1b**). This pattern suggests a shared AAV binding mechanism between microglia and bone marrow macrophages.

**Fig. 1:**
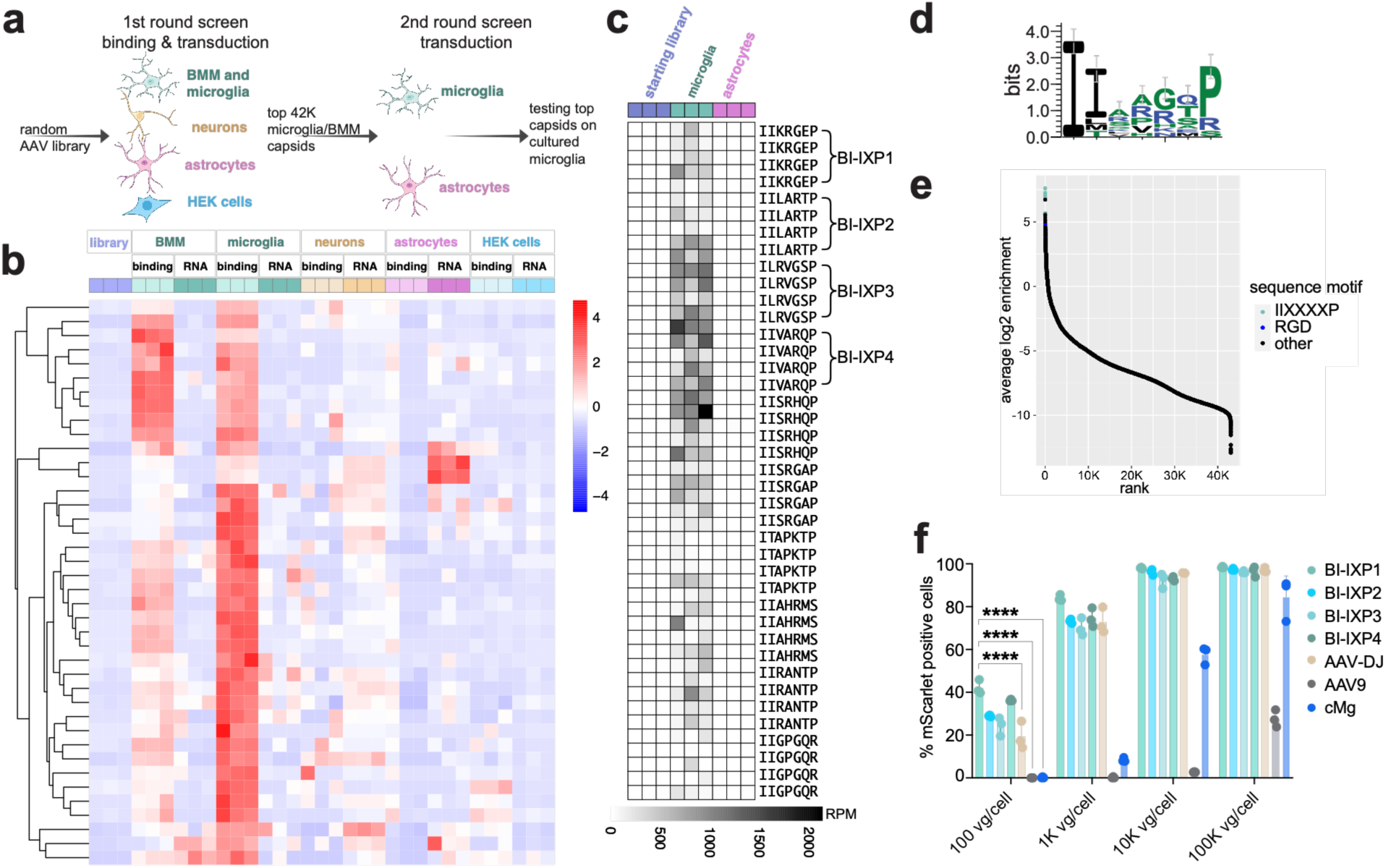
In vitro screening identifies novel capsids that transduce cultured mouse microglia with high efficiency. a, Schematic of the capsid screen. In the first round, AAV9 library with random 7-mer insertions in the capsid protein was added to four cell types followed by two types of assays (binding and transduction). In the second round, top capsids binding to microglia or BMMs were screened for transduction of primary microglia and astrocytes. The top variants were cloned and tested individually for their ability to transduce primary microglia. **b,** Heat map representing top microglia-binding sequences (see methods) in the 1st round screen (colors represent z-scores calculated from RPM, for each row). Columns represent all conditions in the 1st round screen (starting library, binding (labelled ‘binding’) and transduction assays (labelled ‘RNA’, three replicates for each) on 5 different cell types. Each row represents a 7-mer sequence (clustered based on the pattern of abundance in each row). **c,** Heat map representing top ten 7-mer sequences recovered from 2nd round selection on microglia. Each row represents a 7-mer sequence derived from a unique DNA sequence. 7-mer sequences are sorted by average log2 enrichment of 4 codon replicates (mean RPM in microglia vs. starting library), most enriched sequences are at the top. Columns represent samples from the 2nd round selection screen and the input AAV library. **d,** Nucleotide logo of top sequences recovered from the 2nd round screen. **e,** Rank-plot of capsids detected in microglia in the 2nd round screen. Top capsids containing the consensus motif are shown in red. **f,** Transduction of primary mouse microglia with four top capsids recovered from the 2nd round in vitro screen compared to AAV9, AAV-DJ and cMg capsids packaging AAV-CAG-mScarlet-NLS. Transduction was measured by FACS. Points represent individual wells. Error bars represent SD. Ordinary one-way ANOVA (F (8, 18) = 107.9, p-value < 0.0001) with Tukey’s post-hoc multiple-comparisons test was performed to calculate p-values for each condition. Select post-hoc comparisons are shown for 100 vg/cell. **** represents p-value < 0.0001. The 95% confidence intervals for the differences between the means were 33.8% to 50.0% (BI-IXP1 vs. cMg), 34.0% to 50.2% (BI-IXP1 vs. AAV9), and 14.7% to 30.9% (BI-IXP1 vs. AAV-DJ).

We then produced a second-round library consisting of ∼42K unique capsid sequences that either bound to or transduced microglia or BMMs *in vitro*. Each amino acid sequence in the library was encoded by 4 different redundant nucleotide sequences to provide replicates. We performed second-round capsid selection on mouse microglia and astrocytes FACS-purified from mixed culture. Sorting top sequences by the mean enrichment of the 4 replicates revealed a clear consensus motif among top sequences (**Fig. 1c, d and e**). The motif had hydrophobic, branched amino acids at the beginning of the 7-mer insertion (typically two isoleucines), followed by four amino acids of any kind, and typically a proline in the last position (IIXXXXP). We cloned four of the top capsid sequences (named BI-IXP1 through BI-IXP4) and tested them individually on cultured primary microglia by measuring mScarlet reporter gene expression with flow cytometry. The top microglia-targeting capsids transduced a significantly greater fraction of microglia than AAV9 or cMg, a capsid reported to be highly efficient at transducing cultured microglia (Lin et al., 2022) across all tested doses (**Fig. 1f**). We also evaluated microglia transduction by AAV-DJ, as this capsid transduces various cell types with high efficiency (Grimm et al., 2008). AAV-DJ was effective at transducing microglia, but our top capsids were still significantly more efficient at low AAV concentrations (100 vg/cell) (**Fig. 1f**). Finally, we tested the mouse microglia-targeting capsids on human ESC-derived microglia (iMGLs), but we did not detect the same high level of reporter gene expression (**Supplementary Fig. 3**).

### *In vitro* screening identifies AAV capsids that efficiently transduce human iMGLs

Given our success with *in vitro* capsid selection on mouse microglia, we performed a similar capsid selection on human ESC-derived microglia (iMGLs). We recently published a method for lentiviral transduction of iMGLs that utilizes viral-like particles (VLPs) containing Vpx, a protein from simian immunodeficiency virus that improves reverse transcription of the lentiviral genome (Dolan et al., 2023). However, transduction of microglia with lentiviral vectors induced a transient immune response in these cells, characterized by increased expression of interferon stimulated genes. We hypothesized that AAVs might have less of an effect on the transcriptome of iMGLs, given that AAVs are known to be less immunogenic (Nayak & Herzog, 2010). Developing efficient AAV tools for human microglia would provide an important complement to existing lentiviral approaches and enable scalable genetic manipulation of human iMGLs.

The schematic of the selection is shown in **Fig. 2a**. For the first round of selection, we used a capsid library with random 7-mer amino acid insertions in the 588 position of the AAV9 capsid protein. For the binding assay, we added the AAV capsid library to iMGLs after 35 days of *in vitro* differentiation and harvested DNA after incubation at 4°C to collect AAV particles that bound to iMGLs. In addition, we performed a transduction assay, incubating iMGLs with AAV particles for 5 days before collecting RNA. Likely due to the high resistance of iMGLs to AAV transduction, we were unable to recover AAV RNA from the cells. However, we successfully recovered and sequenced AAV DNA from the binding assay.

**Fig. 2:**
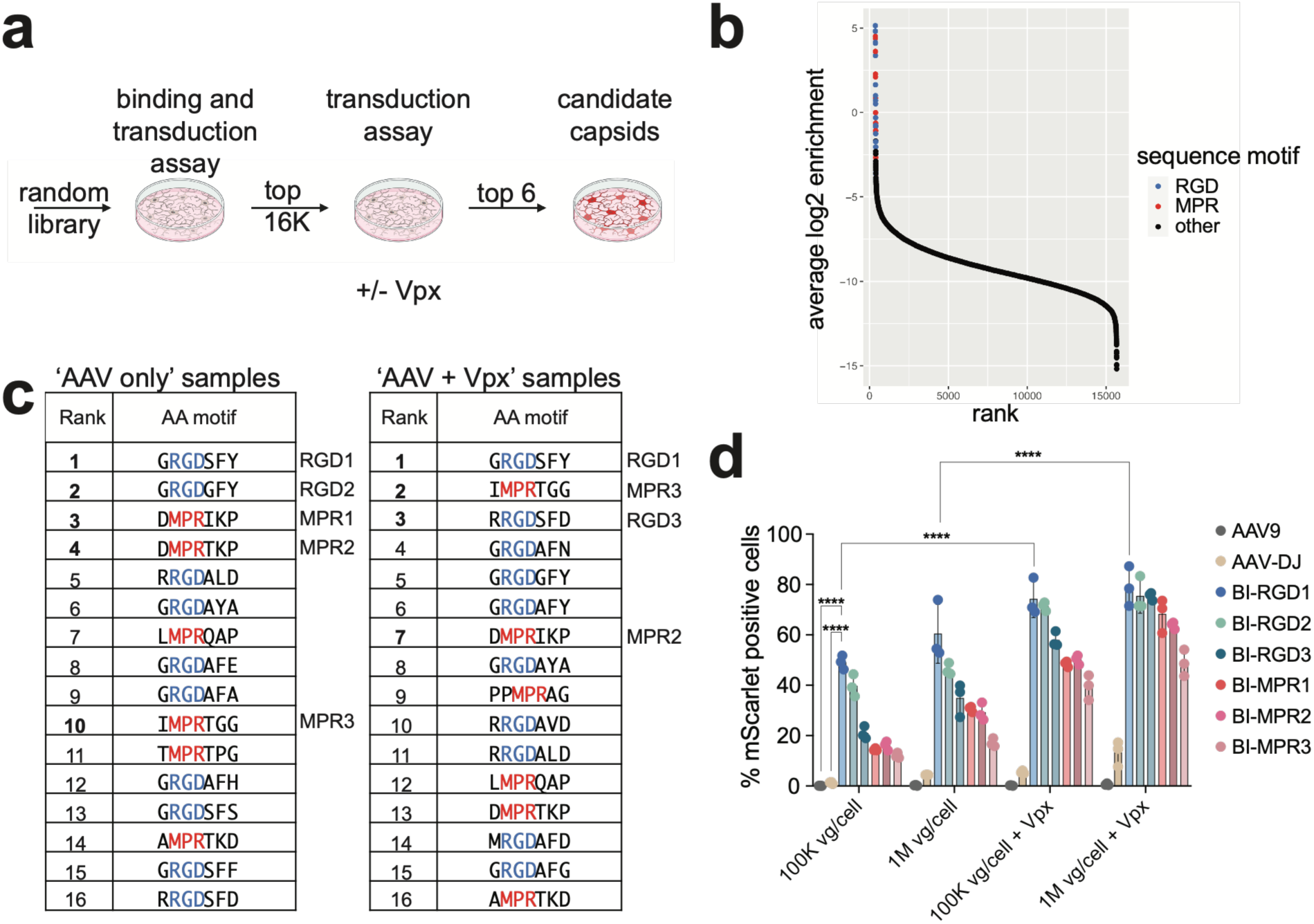
Top capsids from an in vitro screen efficiently transduce iMGLs. a, Schematic of the in vitro capsid screen on iMGLs. **b,** Rank-plot of top capsids from two rounds of capsid screening on iMGLs. Two motifs were identified (shown in blue and red) **c,** Tables of top sequences recovered from the screen. The labels on the right side of the tables show which sequences were cloned for individual testing. **d,** In vitro transduction efficiency of top capsids from the screen. In the +Vpx conditions viral-like particles containing Vpx protein were added along with AAV. Points represent individual wells. Error bars represent SD. Two-way ANOVA (condition: F (3, 64) = 324.3, p-value < 0.0001; capsid: F (7, 64) = 375.7, p-value < 0.0001) with Tukey’s post-hoc multiple-comparisons test was performed. Select post-hoc comparisons are shown between capsids and for BI- RGD1 to demonstrate significantly increased transduction after addition of Vpx viral-like particles. **** represents p-value<0.0001. The 95% confidence intervals for the differences between means were 38.45% to 59.62% (BI-RGD1 vs. AAV9) and 37.28% to 58.45% (BI-RGD1 vs. AAV-DJ) at 100K vg/cell, 16.24% to 34.06% (BI-RGD1 at 100K vg/cell without vs. with Vpx) and 9.71% to 27.53% (BI-RGD1 at 1M vg/cell without vs. with Vpx).

We then made a second-round library of top capsids derived from this first-round binding assay. For the second round of selection, we only performed the transduction assay, given that the second-round library was small and enriched for microglia-binding capsids. In addition to exposing iMGLs to the second-round AAV library alone, we tested whether addition of Vpx-containing VLPs with the AAV library would increase the amount of recovered RNA, as Vpx increases the levels of dNTPs required for second-strand DNA synthesis of AAV. Indeed, when we amplified AAV cDNA from iMGLs there was a clear difference in the amount of AAV cDNA between the two conditions – the samples with Vpx came up ∼2 cycles earlier in qPCR. Sequencing analysis of the most enriched capsids from the second-round transduction assay showed that the top sequences belonged to two consensus clusters (**Fig. 2b**).

One cluster contained an RGD motif at positions 2-4 within the 7-mer (XRGDXXX), which we identified previously in the mouse microglia capsid screen (**Fig. 1d**), but which was not as highly enriched as the IIXXXXP capsids. This motif was previously identified in AAV9 capsids with 7-mer insertions selected for muscle transduction (Tabebordbar et al., 2021; Weinmann et al., 2020). However, unlike the RGD motif in our capsids, the muscle targeting motif is present at positions 1-3 of the 7-mer. The other cluster contained a novel MPR motif (XMPRXXX). We cloned six top capsids containing the two motifs (three for each motif, named BI-RGD1-3 and BI-MPR1-3, **Fig. 2c**) and tested them on iMGLs to assess their transduction efficiency. All top capsids transduced iMGLs with higher efficiency than AAV9 or AAV-DJ. At 100K vg/cell, the top capsid from the screen, BI-RGD1, transduced 49% of iMGLs, whereas AAV9 and AAV-DJ only transduced 0.07% and 1.2% of cells, respectively. We also tested AAV in combination with Vpx VLPs and observed significantly higher transduction when adding Vpx VLPs vs. AAV only (**Fig. 2d**).

Although engineered AAV capsids significantly improved transduction of iMGLs, we observed that transduction led to cell detachment and clumping. When we directly compared cells treated with BI-RGD1 to those treated with the amount of lentivirus that leads to a similar percentage of transduced cells, AAV-treated cells appeared morphologically abnormal (**Supplementary Fig. 4a**). We observed a similar effect when we treated iMGLs with BI-MPR1; however, no morphological changes were seen with AAV9, which does not transduce iMGLs (**Supplementary Fig. 4b**). Sequencing of RNA from AAV-treated iMGLs revealed gene expression changes consistent with an antiviral response and decreased cell cycle progression (**Supplementary Fig. 4c and d**).

Toll-like receptor 9 (TLR9) signaling, induced by the presence of unmethylated CpG dinucleotides in AAV DNA (Hemmi et al., 2000), contributes to the immune response to AAV vectors (Zhu et al., 2009). Given that the CAG promoter present in AAVs used in these experiments contains 98 CpGs, we tested whether an AAV with the human *IBA1* promoter (*hIBA1*) (Serrano et al., 2024), would decrease detachment of iMGLs, as the *hIBA1* promoter contains only a single CpG dinucleotide. Despite significantly reduced expression from the *hIBA1* promoter containing AAVs compared to the CAG containing AAVs, we still observed detached and clumped cells in the culture (**Supplementary Fig. 4d and f**), suggesting that either CpGs in the AAV genome outside of the promoter region trigger the immune response, or that additional mechanisms beyond TLR9 signaling are mediating the response. In summary, while engineered AAVs enable transduction of human iMGLs, they elicit an anti-viral response and morphological changes in these cells that appear to be more pronounced than with lentivirus. Therefore, we conclude that lentivirus is currently the best approach for genetically modifying iMGLs.

### iMGL-targeting capsids are highly efficient at targeting human ESC-derived neurons and astrocytes and can be used unpurified

Due to their broad cellular tropism and strong safety profile, AAVs have emerged as a leading vector for gene therapies targeting neurological disorders (Ling et al., 2023). Patient-derived induced pluripotent stem cell (iPSC) models provide a valuable *in vitro* system for studying disease-relevant human cell types and evaluating candidate AAV- based therapeutic strategies prior to clinical translation. Given the high efficiency of the novel capsids recovered from the iMGL screen, we next tested whether these capsids could also efficiently transduce other human brain cell types. We tested the top capsid from each consensus motif (BI-RGD1 and BI-MPR1) on human ESC-derived neurons and astrocytes. The two capsids were much more efficient at transducing neurons and astrocytes compared to AAV9 when tested at 10K or 100K vg/cell (**Fig. 3a**). Based on total fluorescence per cell, BI-MPR1 was the most efficient capsid for transducing neurons, and BI-RGD1 was the most efficient capsid for transducing astrocytes (**Fig. 3b**). Interestingly, adding Vpx significantly increased transgene expression in neurons treated with BI-MPR1 at 100K vg/cell. In contrast, adding Vpx did not enhance astrocyte transduction (**Fig. 3b**). This observation suggests that dNTPs are limiting for AAV transduction of human ESC-derived neurons.

**Fig. 3:**
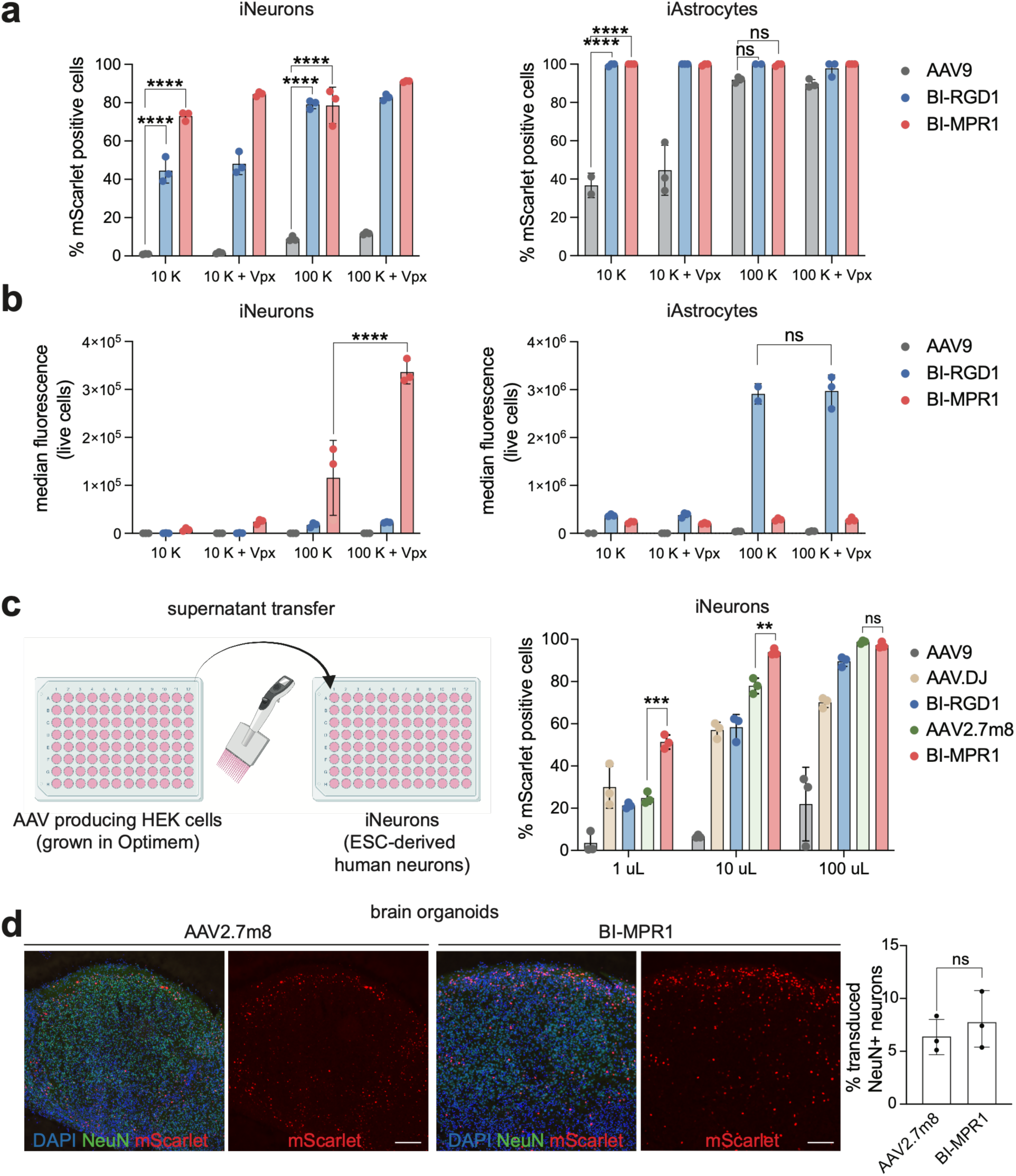
Efficient transduction of human neurons and astrocytes by BI-RGD1 and BI- MPR1. Transduction of human pluripotent stem cells-derived neurons and astrocytes with top capsids recovered from AAV capsid screen on iMGLs compared to AAV9. The AAV genome was CAG-mScarlet-NLS for all experiments. **a,** Percentage of transduced cells. **b,** Median fluorescence of live cells. Transduction was measured by FACS. In **a** and **b** points represent individual wells and error bars represent SD. For each dataset, two-way ANOVA was performed (for iNeurons in **a:** condition: F (3, 24) = 60.61; capsid: F (2, 24) = 1238, for iAstrocytes in **a:** condition: F (3, 22) = 38.29; capsid: F (2, 22) = 213.0, for iNeurons in **b:** condition: F (3, 24) = 46.27; capsid: F (2, 24) = 95.59, p-value for all tests < 0.0001). ANOVA was followed by Tukey’s multiple comparisons test. Select post-hoc comparisons are shown (demonstrating the comparison between AAV9 and the two engineered capsids at two different amounts of virus). In addition, in **b,** comparison between the conditions with and without Vpx is shown, demonstrating an effect of adding Vpx. The 95% confidence intervals for the differences between means for capsid comparisons at 10K vg/cell were 35.6% to 51.5% (BI-RGD1 vs. AAV9) and 64.1% to 80.0% (BI-MPR1 vs. AAV9) on iNeurons and 52.7% to 73.0% (BI-RGD1 vs. AAV9) and 53.2% to 73.5% (BI-MPR1 vs. AAV9) on iAstrocytes. 95% confidence interval for the difference between means of the median fluorescence of live iNeurons with BI-MPR1 without and with Vpx at 100K/vg per cell was 1.6E5 to 2.7E5. **c,** Transduction of human ESC-derived neurons with unpurified AAV9, AAV-DJ, BI-RGD1, AAV2.7m8 and BI-MPR1 capsids. Ordinary one-way ANOVA with Tukey’s multiple comparisons test was performed to calculate p-values for comparisons between different capsids for each volume of AAV containing media (all p-values from ANOVA were < 0.0001). p-values for the comparisons between AAV2.7m8 vs. BI-MPR1 are shown on the graph. 95% confidence intervals for the difference between the means for the comparisons were: 12.2% to 41.0% (for 1 uL), 6.0% to 25.8% (for 10 uL) and -23.0% to 19.9% (for 100 uL). Points in **a-c** represent cells from individual wells and error bars represent SD. **d,** Transduction of NeuN+ neurons in human cortical organoids with AAV2.7m8 and BI- MPR1 capsids. Scale bars = 100 μm. N=3 organoids, 4-8 sections per replicate were quantified. Unpaired, two-tailed Student’s t-test was performed to calculate the p-value (p=0.5013).

Purifying and concentrating AAVs is labor-intensive and outsourcing AAV production can be expensive and slow. We reasoned that highly efficient capsids could be used without purification and concentration for *in vitro* applications. This approach would be especially valuable to simplify applications where a large number of AAVs need to be produced, such as high-throughput arrayed screens. Building on the high transduction efficiency of the BI-MPR1 and BI-RGD1 capsids on human neurons, we tested whether human neurons could be transduced with unpurified and unconcentrated AAV in the form of the media from AAV-producing HEK293T cells. We transfected HEK293T cells in a 96-well plate with plasmids for AAV production, ensuring equal cell numbers and consistent amounts of transfection reagents across all capsids. The next day, we changed media to serum-free Optimem. We collected the media 3 days after transfection and added increasing amounts of media to cultured neurons. In addition to BI-RGD1 and BI-MPR1 capsids, we also tested AAV9 as a control and AAV-DJ and AAV2.7m8 capsids that are known to be highly efficient at transducing various cell types (Grimm et al., 2008; Gurtsieva et al., 2024). The BI-MPR1 capsid was the most efficient from all the capsids we tested, transducing ∼50% of neurons with just 1 uL of media from AAV-producing HEK293T cells (**Fig. 3c**).

The high transduction efficiency of the BI-MPR1 capsid on human ESC-derived neurons also motivated us to test whether this capsid could be useful for transducing human cortical organoids (Velasco et al., 2019). In a recent study that compared a large panel of AAV capsids, AAV2.7m8 was one of the most efficient capsids at transducing brain organoids (Drouyer et al., 2024), therefore we compared AAV2.7m8 side by side with BI-MPR1. Our results show that BI-MPR1 was as efficient as AAV2.7m8 at transducing NeuN+ neurons in cortical organoids (**Fig. 3d**). Importantly, BI-MPR1 can be produced at high titers, whereas AAV2.7m8 yielded a more than sevenfold lower titer in our hands following iodixanol gradient ultracentrifugation, in line with a previous report describing lower production yields for AAV2.7m8 (Cui et al., 2024).

Finally, we tested whether BI-MPR1 could target other immune cell types. We evaluated its efficiency to transduce cultured primary human T cells, a cell type most efficiently targeted by AAV6 (Eyquem et al., 2017; Wang et al., 2016). BI-MPR1 matched transduction efficiency of AAV6 (**Supplementary Fig. 5**) and could serve as an alternative for *in vitro* genome engineering of T cells, particularly given the poor production of AAV6 (Khaparde et al., 2025).

### The efficiency and specificity of microglia targeting *in vivo* are determined by DNA regulatory elements rather than by the capsid

To investigate the influence of capsid identity and viral genome design on the efficiency and specificity of microglial transduction *in vivo*, we performed a systematic comparison of our newly engineered and previously reported microglia-targeting capsids under identical experimental conditions.

We initially tested BI-IXP1, our top capsid from the 2nd round *in vitro* screen, in *vivo* in combination with a CAG-mScarlet-NLS genome. We observed mScarlet expression in non-microglia cells as well as widespread microgliosis (**Supplementary Fig. 6b and d**). We also observed comparable microgliosis with the AAV9 capsid, suggesting that capsid modifications in BI-IXP1 are unlikely to be the cause of the response (**Supplementary Fig. 6a**). To determine whether the microgliosis was driven by reporter expression or by a response to the AAV genome or capsid, we administered AAV9 particles carrying a genome with the CAG promoter but no reporter transgene. Microgliosis was present even in the absence of reporter expression, suggesting that the CAG promoter or other AAV genome element was at least partially responsible for the microgliosis (**Supplementary Fig. 6b and c**).

Having established that the CAG genome was unsuitable for microglial targeting *in vivo*, we next examined recently published genome designs that had been reported to improve microglial transduction. Okada et. al. reported that efficient and relatively specific microglial transduction can be achieved using AAV9 capsid in combination with an AAV genome harboring a mouse *Iba1* promoter and miR-129 and miR-9 target sites (Okada et al., 2022). Luo et al. reported a similar approach using AAV11 capsid, without miR-129 target sites (Luo et al., 2024). The result from Okada et al. was later replicated using an engineered capsid (MC5) which showed significantly higher efficiency compared to AAV9 (Santoscoy et al., 2024). However, both groups reported that the mouse *Iba1* promoter combined with miR-129 and miR-9 target sites resulted in some off-target expression and modest efficiency. Additional engineering of this genome improved its specificity, but its efficiency was still moderate (Aoki et al., 2025). In contrast, Serrano et. al. reported efficient microglial transduction in the striatum (∼86% around the injection site) with near complete specificity using AAV5 in combination with a human *IBA1* (*hIBA1*) promoter and miR-124 target sites (Serrano et al., 2024).

We therefore performed an *in vivo* test of the top capsids from our *in vitro* screen (BI-IXP1, BI-MPR1 and BI-RGD1) combined with a genome published by Serrano et. al. (we replaced GFP in the original genome with mScarlet-NLS, we refer to the genome as hIBA1-mScarlet-miR124TS throughout the manuscript). We compared our capsids with published microglia-targeting capsids: AAV5 (Serrano et al., 2024), MC5 (Santoscoy et al., 2024), AAV11 (Luo et al., 2024), MG1.2 (Lin et al., 2022) and AAV9 as a control. Surprisingly, all the capsids we tested showed comparable, near complete transduction of microglia within the transduction field in both striatum and cortex (**Fig. 4**). The result was unexpected, given that the identity of the capsid had a significant effect on microglia transduction in prior *in vivo* studies and in our *in vitro* experiments. In addition, compared to the genome with the CAG promoter, the genome with *hIBA1* promoter did not cause widespread microgliosis (**Fig. 4a and b and Supplementary Fig. 6b and c**).

**Fig. 4.**
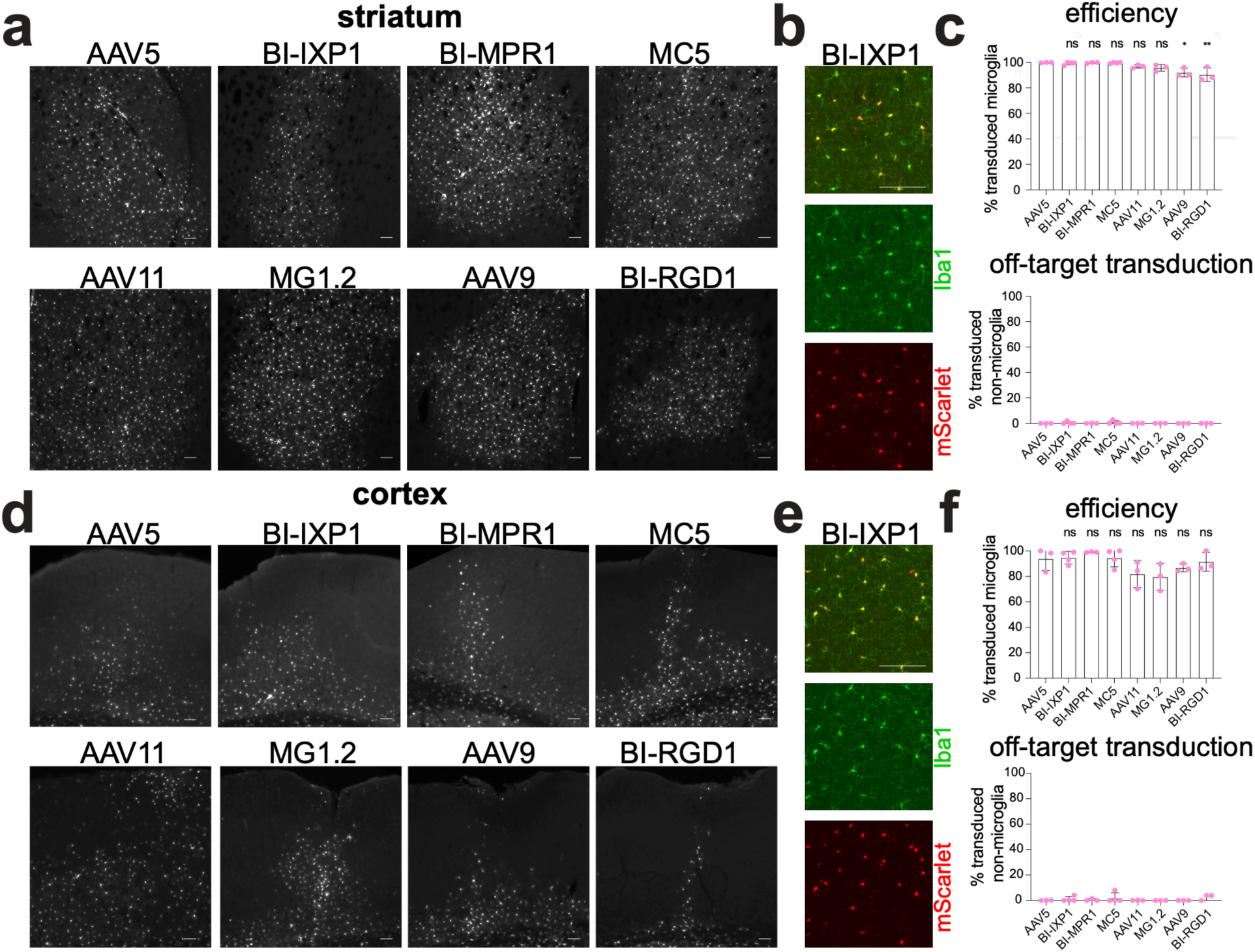
Engineered and natural AAV capsids show comparably high transduction efficiency and specificity after intracranial injection. a, Intrastriatal injections of AAV particles with different capsids (AAV5, BI-IXP1, BI-MPR1, MC5, AAV11, MG1.2, AAV9 and BI-RGD1). All capsids contain the hIBA1-mScarlet-miR124TS genome (modified from Serrano et. al, 2024). **b,** A close-up image of the striatum injected with BI-IXP1 showing overlap between microglia marker Iba1 and mScarlet expressed from the viral genome. **c,** Quantification of transduction efficiency. The top panel shows the percentage of mScarlet+, Iba1+ microglia at the injection site (i.e. efficiency of transduction). The bottom panel shows the percentage of mScarlet positive cells at the injection site that are not microglia (i.e. specificity of transduction). **d-f,** Same analysis as in a-c, but for the cortex. In c and f, points represent individual mice, n=3-4 mice per capsid. Error bars represent SD. Ordinary one-way ANOVA followed by Tukey’s post hoc test was used to calculate p-values. F-statistics and p-values from ANOVA were: F (7, 18) = 7.465, P=0.0003 (transduction efficiency in the striatum); F (7, 18) = 0.6462, P=0.7129 (off-target transduction in the striatum); F (7, 18) = 2.662, P=0.0446 (transduction efficiency in the cortex); F (7, 18) = 0.6192, P=0.7335 (off-target transduction in the cortex). p-values for select post-hoc comparisons between AAV5 and the rest of the capsids are shown above the bar graphs. ns = non-significant p-value, * p < 0.05 and ** p < 0.01. 95% confidence intervals for the differences between mean microglia transduction efficiency in the striatum were -5.5% to 6.9% (AAV5 vs. BI-IXP1), - 6.5% to 6.8% (AAV5 vs. BI-MPR1), -5.9% to 6.5% (AAV5 vs. MC5), -3.6% to 9.7% (AAV5 vs. AAV11), -2.5% to 10.8% (AAV5 vs. MG1.2), 1.1% to 14.4% (AAV5 vs. AAV9), 2.8% to 16% (AAV5 vs. BI-RGD1). In the cortex, they were -19.9% to 18.3% (AAV5 vs. BI-IXP1), - 25.7% to 15.3% (AAV5 vs. BI-MPR1), -19.7% to 18.5% (AAV5 vs. MC5), -8.5% to 32.4% (AAV5 vs. AAV1), -6.3% to 34.7% (AAV5 vs. MG1.2), -13.3% to 27.6% (AAV5 vs. AAV9), -18.1% to 22.8% (AAV5 vs. BI-RGD1). Scale bars in panels a, b, d and e = 100 μm.

Given the efficient transduction observed when using the hIBA*1*-mScarlet- miR124TS AAV genome, we wondered whether we would be able to detect AAV genomes in microglia with *in situ* hybridization. Using RNAscope with an AAV genome- specific probe set, we did not detect the accumulation of AAV genomes inside microglia nuclei. This suggests that AAV internalization and/or trafficking remains a bottleneck preventing viral genomes from efficiently reaching the nucleus, but that transduction is nonetheless achievable with an AAV genome designed for microglia-specific expression (**Supplementary Fig. 7**). Consistent with the RNAscope result, suggesting a low number of viral genomes per microglia, native mScarlet fluorescence in microglia was faint compared to expression normally seen in neurons, and required amplification with an anti-RFP antibody. Taken together, our results show that microglia targeting *in vivo* is highly dependent on the choice of gene regulatory elements, with the capsid variant playing a comparatively minor role.

Given the successful microglia targeting via intracranial injection of AAV with the *hIBA1*-mScarlet-miR124TS genome, we tested whether other routes of administration could be used to transduce microglia more broadly. We tested if intracerebroventricular (ICV) injections of AAV with this genome in P1 pups could be used to target microglia across the brain. We found that ICV injection of BI-IXP1 capsid (1.5E10 vg per pup) resulted in sparse transduction in cortex and hippocampus (**Supplementary Fig. 8**). Achieving more efficient transduction in this context may require a higher dose or further method optimization.

### Microglia can be targeted by intravenously injecting a high dose of BBB-crossing capsids

We then tested whether the *hIBA1*-mScarlet-miR124TS genome can mediate microglia transduction using BBB-crossing capsids AAV-PHP.eB (Chan et al., 2017) and 9P31 (Nonnenmacher et al., 2021). Using a standard dose of AAV that is suitable for transducing neurons and astrocytes (1E11 vg per mouse) led to sparse transduction, so we injected 5E12 vg per mouse. We quantified microglia transduction efficiency in the cortex, striatum and hippocampus. This dose resulted in moderate microglia transduction, with considerable variability across samples and brain regions (we detected about 20 to 50% transduced microglia). (**Fig. 5a and b**). However, we noticed that a large proportion of transduced cells were not microglia but instead appeared to be endothelial cells based on morphology. We stained for the vascular glycocalyx using the tomato lectin (TL) and found that the majority of transduced non-microglia cells in the cortex, striatum and hippocampus were TL-positive. The proportion of non-microglial transduced cells that were TL-positive was lower in samples from mice treated with 9P31 compared to AAV- PHP.eB (**Figure 5c and d**). Our data suggests that BBB-crossing capsids combined with the *hIBA1*-mScarlet-miR124TS genome can transduce a substantial number of microglia, but additional engineering of the genome, perhaps through the introduction of additional endothelial-specific miR target sites, will be required to improve the specificity of this approach.

**Fig. 5.**
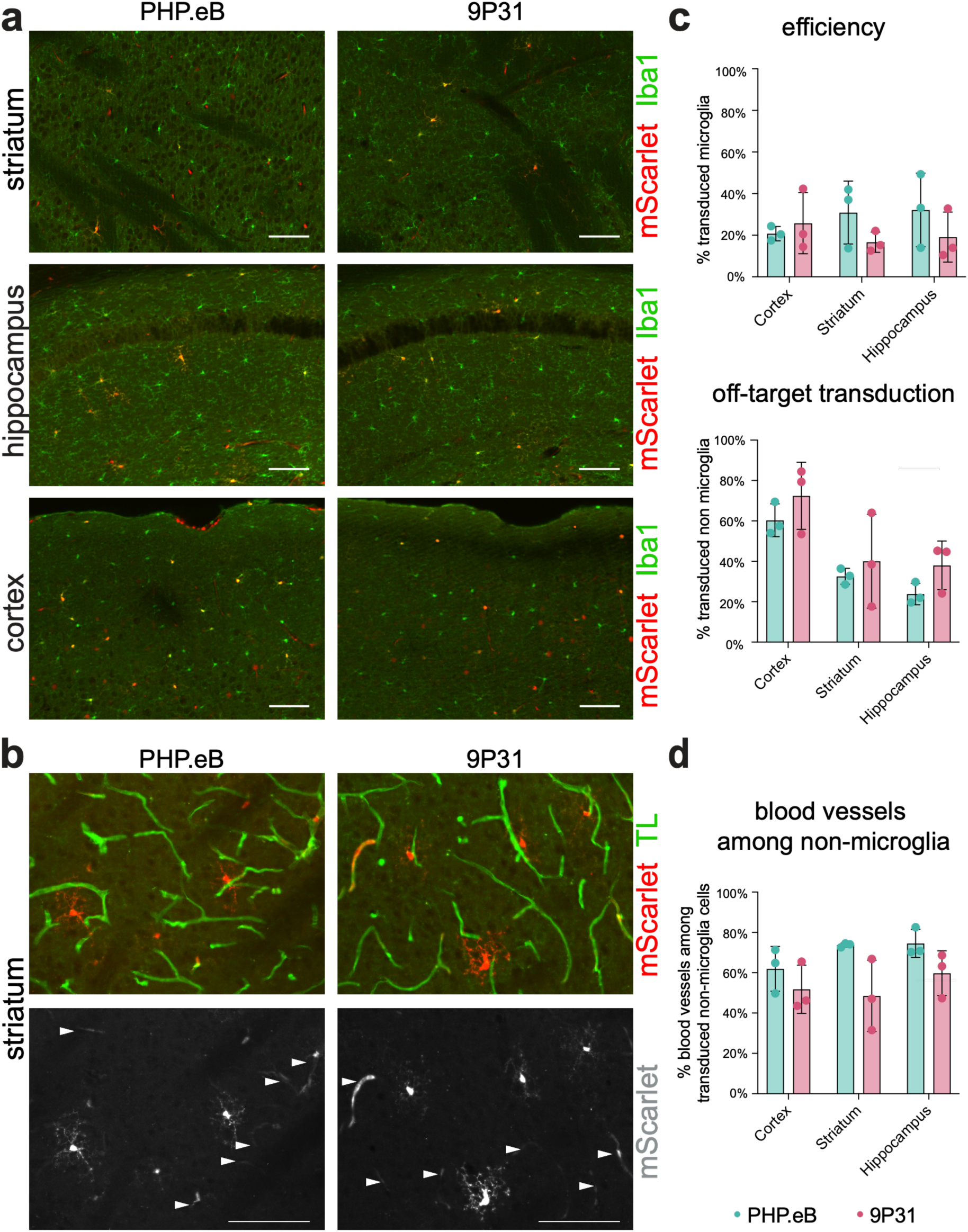
Comparison of microglia transduction efficiency of BBB-crossing capsids containing viral genome with hIBA1 promoter-driven transgene and miR-124 target sites. a, Expression of reporter gene (mScarlet) from intravenously injected BBB-crossing AAV (AAV-PHP.eB and 9P31) in three brain regions: striatum, hippocampus, and cortex. Both capsids contain the hIBA1-mScarlet-miR124TS genome, originally published by Serrano et. al, 2024. Microglia stained with anti-Iba1 antibodies are shown in green. Scale bars = 100 μm. **b,** Analysis of transduction of blood vessels (stained with tomato lectin) in the striatum with the microglia targeting BBB-crossing AAVs. Arrowheads show transduced cells that overlap with blood vessel staining. Scale bars = 100 μm. **c,** Quantification of the microscopy data. The graph on the top shows the percentage of mScarlet positive microglia (determined by Iba1 expression) in different brain regions (i.e. efficiency of transduction). The graph on the bottom shows the percentage of mScarlet positive microglia at the injection site that are not microglia (i.e. specificity of transduction). **d,** Quantification of transduced blood vessels in different brain regions with the two capsids. In **c** and **d,** points represent individual mice, n=3 mice per capsid. Error bars represent SD. Two-tailed Student’s t-tests were used to calculate p-values comparing the two capsids within each brain region. None of the comparisons were statistically significant.

## Discussion

In this study, we evaluated the relative contributions of capsid engineering and viral genome design to efficient microglial transduction and developed new tools for genetic manipulation of mouse and human microglia. We first performed capsid screens on cultured mouse and human ESC-derived microglia and showed that capsid engineering can enhance microglial transduction *in vitro*. These *in vitro* binding and transduction screens establish a generalizable strategy for engineering AAV capsids to target cells that are resistant to viral transduction. Incorporating the binding screen yielded robust, reproducible data in the first round, which was then followed by a second-round transduction assay to identify the most efficient capsids. This two-step approach was particularly important for iMGLs, where initial transduction screens failed to yield signal, but a second-round transduction screen of variants from a first-round binding screen enabled identification of efficient variants. Using this screening strategy, we identified three novel capsids (BI-IXP1, BI-RGD1 and BI-MPR1) that exhibit markedly enhanced transduction efficiency in microglia and additional cell types *in vitro*. Notably, BI-IXP1 outperformed all other capsids in primary mouse microglia, whereas BI-MPR1 showed superior transduction of human ESC-derived neurons. We further showed that BI-MPR1 can be used unpurified on human ESC-derived neurons, facilitating scalable applications such as arrayed CRISPR and ORF screens. Together, these capsids provide broadly useful reagents for genetic manipulation of cultured microglia and other human cell types.

We also observed that iMGLs change morphology and express interferon-stimulated genes upon exposure to AAV. Whether this response reflects an intrinsic sensitivity of human microglia to AAV or is specific to iMGL differentiation or culture conditions remains unclear. Future studies with cultured primary human microglia or with iMGLs in the context of brain environment (Hasselmann et al., 2019) will help clarify this question. If the immune response to AAV is indeed an intrinsic property of human microglia, uncovering the exact mechanisms underlying this antiviral response will enable further optimization of AAV transduction approaches and facilitate the development of AAV-based strategies for microglia-targeted gene therapy. These findings also highlight the importance of carefully evaluating the biological consequences of viral transduction when developing gene delivery approaches for human microglia.

A key finding of this study is the demonstration that viral genome design, rather than capsid identity, is the dominant determinant of efficient and specific microglial transduction *in vivo.* Following intracranial injection in the striatum and cortex, all tested capsids exhibited high levels of microglial transduction and specificity when paired with an optimized genome carrying the *hIBA1* promoter and miR-124 target sites originally reported by Serrano et al. This result contrasts with prior reports suggesting superior performance of specific capsids. While MC5, AAV11 and AAV5 have been reported to outperform AAV9 for microglial targeting (Santoscoy et al. 2024; Luo et al. 2024; Serrano et al. 2024), we observed only modest differences under matched conditions. Importantly, because these previous studies each used different viral genomes for their capsid comparisons, the relative contributions of genome design and capsid engineering could not be distinguished.

When paired with a genome containing *hIBA1* promoter and miR124 target sites all capsids converged on similarly high transduction efficiency and specificity, demonstrating that optimized genome design can override differences in capsid performance *in vivo*. On the other hand, when we tested the AAV9 and BI-IXP1 capsids carrying a genome driven by the CAG promoter, we did not observe the widespread microglia transduction seen with the *hIBA1* promoter. Another group has recently reported a similar observation. When they injected AAV9 carrying mouse *Iba1* promoter in combination with miR-9 and miR-129 target sites in the mouse cortex they observed transduced microglia. However, when mouse *Iba1* promoter was replaced with CAG promoter, they did not see any microglia transduction (Aoki et al., 2025).

In addition, we also observed significant microglia activation when using the CAG promoter *in vivo*, compared to the *hIBA1* promoter. This could be due to virus sensing through TLR9 given that the CAG promoter contains 98 CpGs that activate TLR9 signaling and the *hIBA1* promoter only contains one CpG. The rest of the viral genome we used in our study contains 155 CpGs, so the CpGs in the CAG promoter significantly increase the total number of CpGs in the viral DNA. These observations suggest that viral genome architecture influences not only transgene specificity but also the immune response of microglia to AAV.

Finally, our results highlight limitations of BBB-crossing AAV capsids for microglia targeting. Systemic delivery of BBB-crossing variants packaging the genome with *hIBA1* promoter and miR-124 target sites achieved moderate microglial transduction, but cell-type specificity remained a major challenge. We observed substantial off-target transduction, particularly in vascular endothelial cells, consistent with prior work (Aoki et al., 2025). Thus, further optimization of genome design presents an opportunity to achieve high microglial specificity following vascular transcytosis.

Altogether, our study provides tools for efficient and scalable transduction of microglia and other cell types *in vitro* and identifies the current optimal approach for transducing mouse microglia *in vivo*, as well as remaining challenges. Understanding the optimal approaches to targeting microglia will facilitate studies linking genetic variants and transcriptional states to cellular function in these cells, which play key roles in brain development and disease, and support the development of microglia-directed gene therapies for neurological disease.

## Methods

### Animals

All procedures were performed as approved by the Broad Institute Institutional Animal Care and Use Committee (IACUC). Cx3cr1-Cre-ERT2 mice were purchased from the Jackson Laboratory (strain #021160). WT mice (strain C57BL/6NCr) for mixed culture were purchased from Charles River and from JAX for all other experiments (strain #000664). Mice were euthanized using CO2 inhalation (for flow cytometry and *in vivo* capsid selection using CREATE) or vascular perfusion with PBS and 4% paraformaldehyde in PBS (for RNAscope and characterization of *in vivo* AAV transduction by immunofluorescence staining).

### In vivo AAV administration and tissue processing

Intravenous administration of AAV was performed by anesthetizing adult mice (2% isoflurane and 1 L/min oxygen) and injecting ∼100 µL (depending on the exact titer) of AAV into the retro-orbital sinus (1E12 vg/mouse for PHP.B injections in Supplementary Fig. 1 and 5E12 for injections in Fig. 5).

For *in vivo* capsid selection and testing candidate capsids, AAVs were injected intracranially into the adult mice. Briefly, mice were anesthetized using isoflurane and placed in a stereotaxic frame. A small craniotomy was performed above the injection site (coordinates from bregma for striatum: anteroposterior +0.62 mm, mediolateral ±1.5 mm, dorsoventral -3.0 mm). For capsid selection AAV library was injected in the cortex, in addition to the striatum (dorsoventral coordinate: -0.8 mm). AAV (4E9 vg/injection site for testing candidate capsids and 1E10 vg for the *in vivo* library selection) were loaded into a glass capillary needle and injected at a rate of 1 nL/s using a microinjection pump (Nanoject III, Drummond Scientific Company). After each injection, the needle was left in place for an additional 5-10 minutes to minimize backflow, then slowly withdrawn, and the skin over the craniotomy was sutured.

For ICV injections, 1 uL of BI-IXP1: *hIBA1*-mScarlet-miR124TS (concentration 7.78E12 vg/mL) was injected per hemisphere into P1 pups.

For experiments shown in Fig. 4, Fig 5. and Supplementary Fig. 6 and 8, mice were perfused 3 weeks after injection with 4% PFA in PBS. After perfusion brains were harvested, fixed in 4% PFA in PBS overnight, sectioned using a vibratome to 40 micrometer sections and stained with goat anti-Iba1 antibody (WAKO, 011-27991) 1:500 dilution and rabbit anti-RFP antibody (Rockland, 600-401-379) at 1:100 dilution to amplify the mScarlet signal. To label vasculature in Fig. 5b, fluorophore-conjugated tomato lectin (Thermo Fisher, L32470) at 1:50 dilution was used in addition to the anti-Iba1 and anti-RFP antibodies. All experiments shown in Fig. 4 were performed with investigators blinded to the identity of the AAV capsid injected into each mouse. Blinding was maintained until completion of the data analysis.

### FACS

Cell sorting was performed to purify astrocytes and microglia from mice injected with PHP.B and controls. Mice were perfused with 1X HBSS. Brains were processed using Adult Brain Dissociation Kit (Miltenyi Biotec, 130-107-677) according to the manufacturer’s instructions. Samples were first treated with anti-mouse CD16/CD32 (Fc Block, BD Biosciences, 553142) and then stained with the following antibodies against mouse antigens from Biolegend: anti-CD45 (103113), anti-CD11b (101241), anti-P2RY12 (848005), anti-F4/80 (123133) and anti-ACSA-2 (Miltenyi Biotec, 130-116-243). Microglia were defined as CD45+, CD11b+, P2RY12+ and F4/80+. Astrocytes were negative for microglia markers and positive for ACSA-2.

Cell sorting was also performed to purify microglia from mixed culture to remove astrocytes. As above, samples were first treated with Fc Block and then stained with antibodies against mouse CD45 and CD11b. CD45+ and CD11b+ cells were used for the *in vitro* binding and transduction screens.

### PCR on AAV DNA from microglia and astrocytes

After sorting, AAV DNA was purified from microglia and astrocytes using QIAamp DNA Micro Kit (Qiagen, 56304) for purification of DNA from small samples. qPCR was performed using LightCycler 480 SYBR Green I Master mix (Roche, 04707516001) with primers binding to WPRE in the AAV genome. To normalize the amount of AAV DNA to the amount of sample, we used primers specific to a gene in the mouse genome (mGluco).

### RNAscope

To detect AAV genomes, we used RNAscope Multiplex Fluorescent Reagent Kit v2 (Bio-Techne, 323100) following manufacturer’s instructions. CMV enhancer (559071) and WPRE (582691) probe sets were used to stain AAV genomes. Rabbit anti-Iba1 antibody was used to detect microglia (Biocare, 290) at 1:500 dilution. AAV genomes were detected in the far-red channel using the Cy5 dye (Perkin Elmer, NEL745001KT). Microscopy images were quantified using CellProfiler (Stirling et al., 2021).

### AAV production and titering

AAVs were generated by triple transfection of HEK293T/17 cells (ATCC, CRL-11268) using polyethylenimine (PEI), purified by ultracentrifugation over iodixanol gradients, and titered as previously described in Krolak et. al., 2022.

### Mixed glial cell culture

Mixed glial cell culture, from which microglia were later purified for the *in vitro* capsid screens, were set up as follows: anesthetized P4 pups were decapitated, and their brains were removed into chilled PBS. Meninges were removed under dissection microscope, and tissue was triturated using a 5 mL pipette followed by a glass Pasteur pipette. Cells were then spun down and supernatant was removed. Cells were resuspended in DMEM media and filtered through at 70 um Corning cell strainer (Millipore Sigma, 431751). Cells were then seeded in 175 cm2 flasks (2 brains per flask) in 30 mL DMEM with 10% FBS and incubated at 37°C in the tissue culture incubator. The next day, the media was removed, cells were washed with 10 mL PBS. 60 mL DMEM media with 10% FBS was added to the cells. They were incubated for 13 days in a tissue culture incubator. After 13 days, flasks were shaken overnight on a shaker in a tissue culture incubator. Supernatant was removed to collect microglia. Cells were then spun down, resuspended in fresh media with FBS and stained for FACS with anti-CD45 and anti-CD11b antibodies as described above. Astrocytes were removed from the flask using TrypLE trypsin (Thermo Fisher, 12604021), spun down and resuspended in fresh DMEM media, followed by staining for the ACSA-2 antigen and FACS sorting.

### Library design and cloning

For transduction and binding assays, we used an AAV genome vector that was designed to enrich for functional AAV capsid sequences by recovering capsid mRNA from transduced cells (described in more detail in Krolak et. al., 2022). Briefly, the vector uses a ubiquitous CMV enhancer and AAV5 p41 gene regulatory elements to drive AAV9 Cap expression. A previously described cap-deficient Rep-AAP AAV helper plasmid (Deverman et. al., 2016) was supplied in trans to generate AAV. The initial random 7-mer library was produced using 5’-CGGACTCAGACTATCAGCTCCC-3’ and 5’- GTATTCCTTGGTTTTGAACCCAACCGGTCTGCGCCTGTGCMNNMNNMNNMNNMNNMNNMNNTTGGGCACTCTGGTGGTTTGTG-3’ primers (IDT) to PCR amplify a modified AAV9 template (K449R) using Q5 Hot Start High-Fidelity 2X Master Mix (NEB, M0494S) following the manufacturer’s protocol. Note that the second primer contains MNN nucleotides, which correspond to NNK in the forward direction (K in the third position is used to decrease the frequency of stop codons in the library). The PCR products were run on agarose gel followed by purification using Qiaquick Gel Extraction kit (Qiagen, 28704). The PCR insert was assembled into a linearized mRNA selection vector with NEBuilder HiFi DNA Assembly Master Mix (NEB, E2621L). Afterwards, Quick CIP (NEB, M0508S) was spiked into the reaction and incubated at 37°C for 30 minutes to dephosphorylate unincorporated dNTPs. Finally, T5 Exonuclease (NEB M0663S) was added to the reaction mixture and incubated at 37°C for 30 minutes to remove unassembled products. The final assembled product was cleaned up using AMPure XP beads (Beckman, A63881) following the manufacturer’s protocol and quantified using the Qubit dsDNA HS Assay Kit (Thermo Fisher, Q32851). Capsid variants chosen for secondary screening were synthesized as a pool of reverse primers (Agilent). Each capsid 7-mer was represented by four unique nucleotide sequences (called ‘codon replicates’) and cloned into the backbone as described above.

CREATE library was designed as described previously (Deverman et. al., 2016) and assembled using the protocol described above.

### CREATE

The AAV library was injected intracranially as described above. After 3 weeks, mice were euthanized and brain tissue samples were collected. AAV DNA was purified using TRIzol (Thermo Fisher, 15596018) following manufacturer’s instructions for RNA isolation (a significant amount of AAV DNA is found in the aqueous phase (Deverman et. al., 2016)).

### Binding and transduction assays

For the binding and transduction assays to select for mouse microglia-targeting capsids, we used primary mouse microglia and astrocytes from mixed glial culture which we purified as described above. In addition, for the first-round library selection on mouse cells we also used the HEK 293T cells, primary mouse cortical neurons (Thermo Fisher, A15586) and bone marrow macrophages from Balb/C mice (Charles River, 1118- 4983DE20, currently not available anymore). For the binding assay, we exposed cells to the first-round AAV capsid library for 2 hours at 4°C on a rocker to select for capsids with increased binding to microglia and other cell types. For the transduction assay, we exposed the cells to the first and second round AAV library for 5 days at 37°C to select for capsids that successfully transduce microglia and other cell types. For the selection on iMGLs, we added 10 uL of Vpx viral-like particles per 10,000 cells for the Vpx+ condition for second round library selection.

For the binding assay, cells were treated with TrypLE (Thermo Fisher, 12604021), collected in tubes and spun down. DNA from binding assays was purified using the DNEasy blood and tissue kit (Qiagen, 69504). For the transduction assays, cells were lysed directly in the tissue culture dishes using RNA RLT buffer from the RNEasy Plus kit (Qiagen, 74134). RNA was purified following the manufacturer’s instructions. RNA was treated with Turbo DNAse (Thermo Fisher, AM2239) and reverse transcribed using Maxima RT reverse transcriptase (Thermo Fisher, EP0741).

### Sample preparation for sequencing of viral DNA

To prepare AAV libraries for sequencing, qPCR amplification using primer pairs described in Huang et al., 2023 was used to attach partial Illumina Read 1 and Read 2 sequences to AAV DNA and cDNA. Q5 Hot Start High-Fidelity 2X Master Mix (New England Biolabs, M0494L) with added Sybr Green dye (Thermo Fisher, S7567) was used. When fluorescence reached ∼1000 RFUs, samples were taken out of the PCR machine to prevent overamplification. PCR products were run on agarose gel to confirm that they are the right size and purified using QIAquick Gel Extraction Kit (Qiagen, 28704) followed by purification with QIAquick PCR Purification Kit (Qiagen, 28104) to remove any impurities. Then, 1 μL of 10X diluted PCR product was used as input in a second round of PCR to attach on Illumina adapters and dual index primers (NEB, E7600S) for 5-8 PCR cycles using Q5 Hot Start-High-Fidelity 2X Master Mix with an annealing temperature of 65 °C for 20 s and an extension time of 60 s. The round two PCR products were purified using AMPure XP beads following the manufacturer’s protocol and eluted in 25 μL UltraPure DNase/RNase-Free distilled water (Thermo Fisher, 10977023). Eluted DNA was sequenced on MiSeq (Illumina) following manufacturer’s instructions.

### AAV capsid library selection data analysis

Raw sequencing data was processed as described in Eid et. al., 2024. Read counts were normalized to sequencing depth and sorted by average log2 enrichment across codon replicates (which was calculated by first calculating the log2 enrichment vs. starting library for each nucleotide sequence, followed by calculating the mean of the 4 values for each codon replicate). The nucleotide sequence logo shown in Fig. 1d was generated using WebLogo (Crooks et al., 2004).

In the heat map in Fig. 1b, top microglia binding sequences are shown (sorted by RPM) that also have the ratio between mean RPM in microglia / mean RPM in non-macrophage samples higher than 4 (to show sequences that are most specific to microglia).

### Differentiation of human ESC-derived microglia, neurons and astrocytes

H1 human ESC line (from WiCell) was used to generate the differentiated cells. iMGLs were differentiated as previously described (Dolan et al., 2023). To make induced human neurons, we first generated AAVS1 safe harbor NGN2 inducible human ESC line with zeocin resistance conferred in the presence of doxycycline and then differentiated them into excitatory NGN2 neurons (Nehme et al., 2018). Astrocytes were differentiated using AAVS1 safe harbor NFIB-SOX9 inducible human ESC line and a previously described differentiation protocol (Morshed et al., 2026).

### Vpx VLP production

Vpx particles were produced as described previously (Dolan et al., 2023).

### Sequencing of RNA from iMGLs

iMGLs were seeded in 24-well plates on differentiation day 30 and treated on day 35 with either AAV, AAV plus Vpx particles, lentivirus plus Vpx particles, or Vpx particles alone. 5 days later, cells were lysed directly in the tissue culture dishes using RNA RLT buffer from the RNEasy Plus kit (Qiagen, 74134). RNA was purified following the manufacturer’s instructions. RNA was treated with Turbo DNAse (Thermo Fisher, AM2239). Library preparation and sequencing were performed by Novogene Corporation Inc. (Sacramento, CA). Raw sequencing data were aligned using STAR (Dobin et al., 2013), reads mapping to genes were quantified with featureCounts (Liao et al., 2014), and differential expression analysis was performed using DESeq2 (Love et al., 2014). Gene ontology analysis was performed using Enrichr (Xie et al., 2021).

### Differentiation of human induced pluripotent stem cells (iPSC) into cortical organoids

11a human iPSC line, obtained from the Harvard Stem Cell Institute, was used to generate cortical organoids as previously described (Velasco et al., 2019). Cortical organoids were cultured for 4 months and transduced with 50 vg/cell of AAV2.7m8 and BI-MPR1 capsids containing CAG-mScarlet-NLS genome. Individual organoids were incubated with the virus for 24h in Cortical Differentiation Medium IV (CDMIV, without Matrigel) in a 24-well plate containing 500uL of medium per well. The medium was fully exchanged the next day and subsequently replaced once per week. After 14 days of transduction, organoids were fixed in 4% PFA and sectioned using a cryostat. Organoid sections were then mounted on slides and stained with anti-NeuN antibody (AbCam, ab104225) at 1:500 dilution.

### Immunohistochemistry data analysis

Data in Fig. 3d and Supplementary Fig. 6 was analyzed using CellProfiler (Stirling et al., 2021). Briefly, for organoid data analysis, NeuN+ cells were identified using IdentifyPrimaryObjects module. Threshold module was used to identify cells transduced with AAV. MaskObjects module was used to identify cells that are both NeuN+ and AAV+. Data in Fig. 4 and Fig. 5 was analyzed using QuPath (Bankhead et al., 2017). Coronal (Fig. 4, direct injection of AAV) and sagittal (Fig. 5, intravenous injection of AAV) brain sections were analyzed to quantify capsid transduction efficiency and off-target transduction across brain regions. Regions of interest were manually delineated, including cortex, striatum and hippocampus. For on-target transduction efficiency, microglia were identified by cell detection in the green fluorescence channel. A single-threshold classifier was applied to the red channel (AAV-encoded mScarlet), and cells above threshold were classified as transduced microglia. Transduction efficiency was calculated as the percentage of microglia that were mScarlet-positive within each region of interest. For off-target analysis, transduced cells were first detected in the red channel. A single-threshold classifier was then applied to the green channel to identify microglia. Off-target transduction was quantified as the percentage of transduced cells that were not microglia within each region of interest. To quantify vascular endothelial cell transduction, cells detected in the red channel were classified as transduced, and cells above threshold in the green channel were classified as vascular endothelial cells. Transduction was quantified as the percentage of transduced vascular structures within each region of interest using a composite classifier.

## Acknowledgements

We thank Alec Walker for his assistance with mouse microglia FACS and Alexander Svanbergsson, Fatma-Elzahraa Eid, and Albert Chen for their assistance with data analysis. We also thank Shata Dasgupta and Esteban Miglietta for their advice on image analysis with CellProfiler and Jordan Doman for technical assistance with experiments using iMGLs.

Illustrations in Supplementary Fig. 1, Fig 1, Fig. 2 and Fig. 3 were created using BioRender.

This work was supported by grants from MassCATS (to B.D. and B.S.), NIMH (R21 MH126409 to B.D. and B.S.), Stanley Family Foundation (to B.D. and B.S.), Alzheimer’s Association (grants ADSF-23-1054204-C and ADSF-25-1460070-C to B.S.) and the Howard Hughes Medical Institute (to B.S.).

---

**Supplementary Fig. 1:**
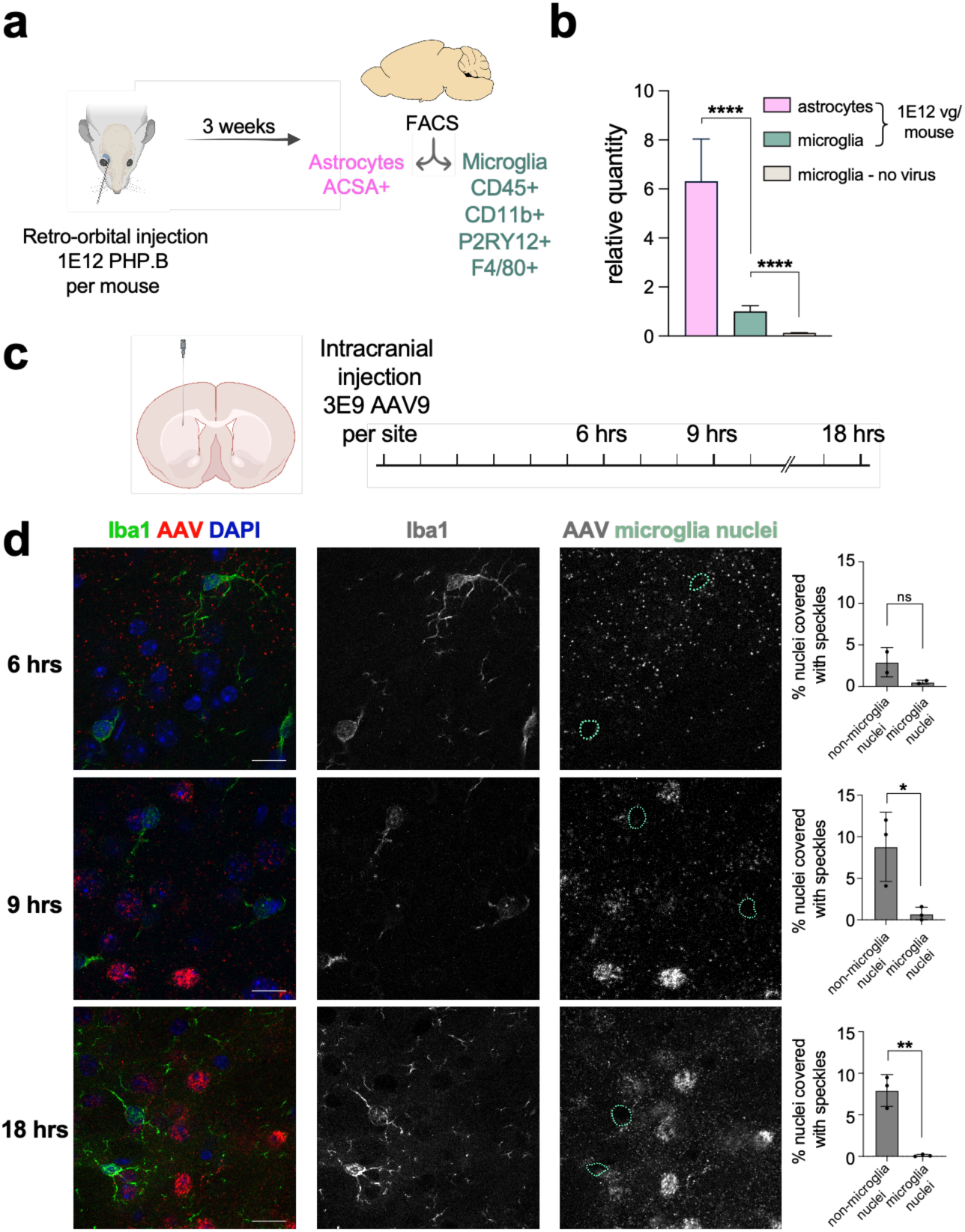
AAV genomes preferentially accumulate in non-microglial cells after intravenous and intracranial injection. a, Schematic of the experiment: AAV was injected intravenously, microglia and astrocytes were FACS sorted 3 weeks after injection, viral DNA was quantified using qPCR. **b,** qPCR amplification of AAV DNA from sorted microglia and astrocytes 3 weeks after intravenous PHP.B injection. The bar chart represents the output of the Bio-Rad CFX Maestro software, which was used to calculate relative AAV DNA quantities and p-values (using an unpaired, two-tailed Student’s t-test). Error bars represent SD. **** represent p < 0.001. n=3 mice. **c,** Schematic of the time course experiment. **d,** In situ hybridization for AAV genomes at 6, 9 and 18 hours after intracranial injection (red, detected by RNAscope, speckles represent internalized AAV genomes). Microglia marker Iba1 is shown in green, and DAPI, a nuclear marker, is shown in blue. In the third column, microglia nuclei defined by DAPI staining are outlined in green. n=3 mice per time point (for 9 hrs and 18 hrs). Scale bars = 20 μm. p-value was calculated using unpaired, two-tailed Student’s t-test. Error bars represent SD. * represents p < 0.05 and ** represents p < 0.01.

**Supplementary Fig. 2:**
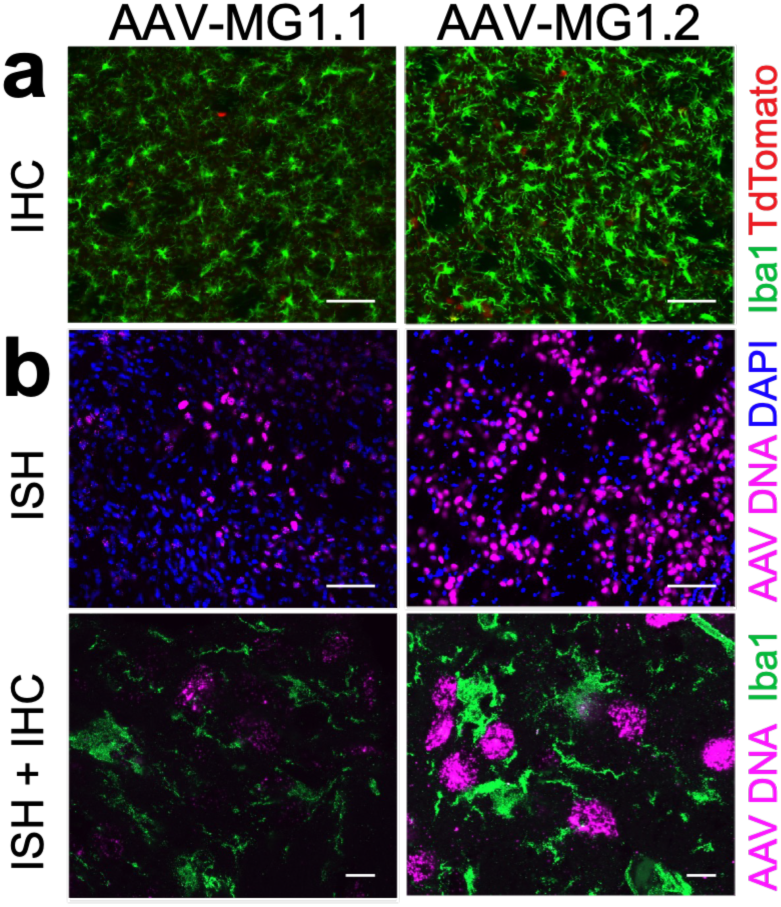
AAV genomes preferentially accumulate in non-microglial cells after intracranial injection of AAV-MG1.1 and AAV-MG1.2. a, Visualization of the injection sites in the striatum. 4E9 vg/site of MG1.1 and MG1.2 packaged with (Cre- dependent) SFFV-DIO-TdTomato genome were injected into Cx3cr1-Cre-ER mice that express Cre in microglia in a tamoxifen-dependent manner. 4.5 mg of tamoxifen was injected intraperitoneally starting 1 week after AAV injection (spread over 3 days), tissue was harvested 3 weeks after AAV injection. Microglia were stained with a microglia marker Iba1. Scale bars = 100 μm. **b,** In situ hybridization for AAV genomes after intracranial injection (purple, detected by RNAscope). Microglia marker Iba1 is shown in green, and DAPI, a nuclear marker, is shown in blue. n=3 mice per capsid. Scale bars = 100 μm (top panels) and 10 μm (bottom panels).

**Supplementary Fig. 3:**
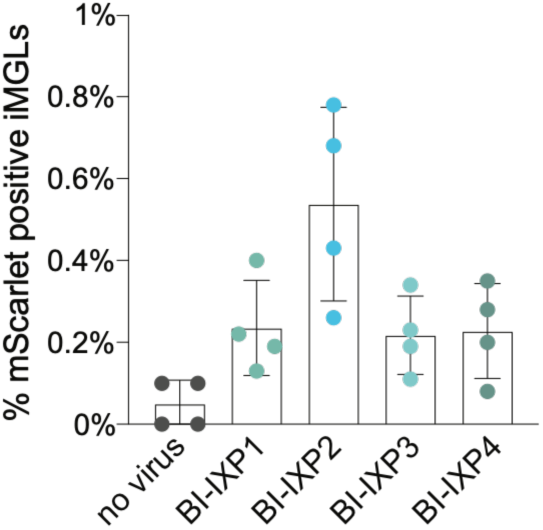
Engineered mouse microglia-targeting capsids do not transduce human microglia (iMGLs) with the same efficiency as mouse microglia. Transduction of human iMGLs with top four capsids recovered from the 2nd round in vitro screen packaging AAV-CAG-mScarlet-NLS genome. iMGLs were treated with 10,000 vg/cell. Transduction was measured by FACS 5 days after AAVs were added to the cells.

**Supplementary Fig. 4:**
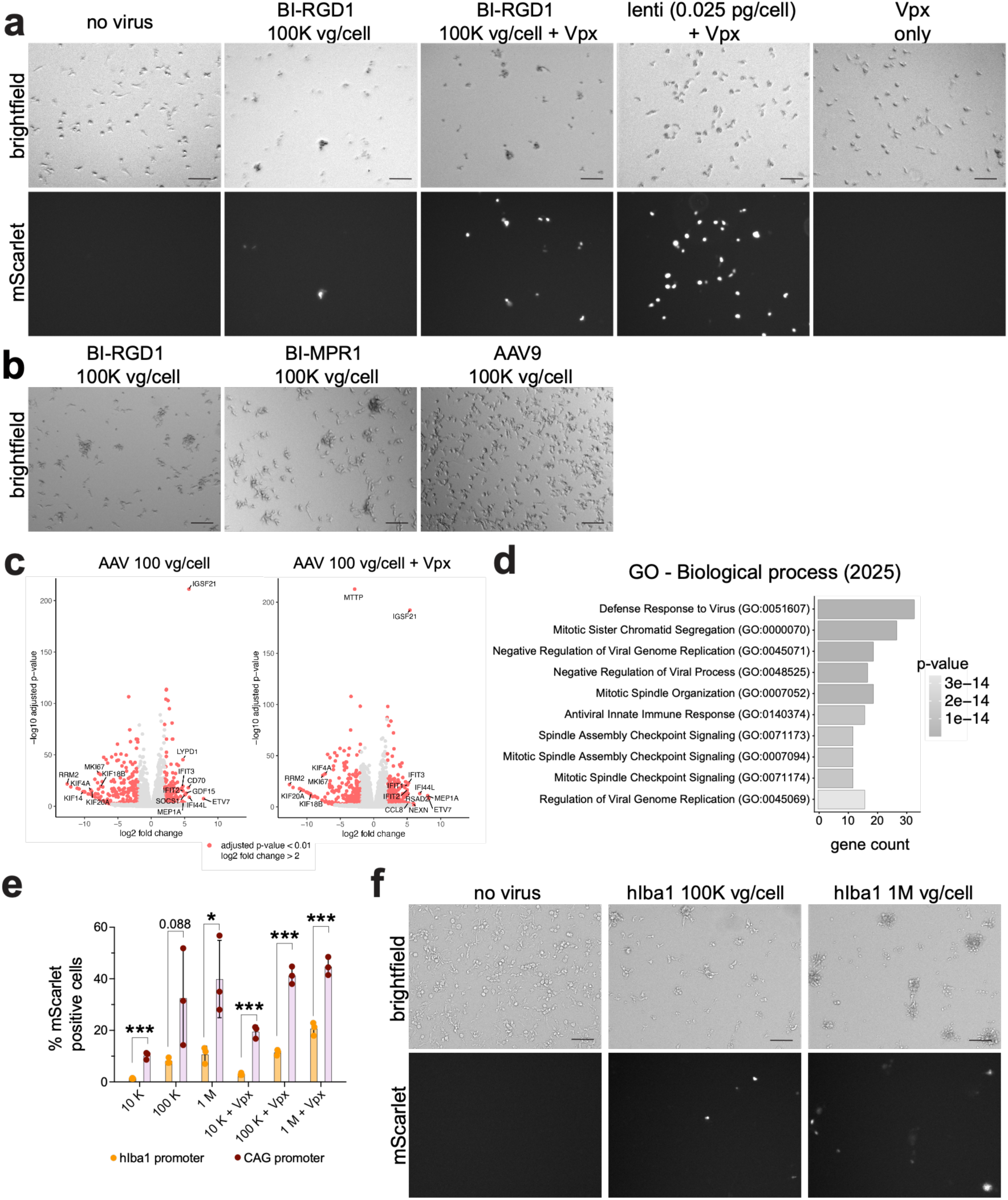
AAV treatment induces cell detachment and antiviral response in iMGLs. a, Microscopy images showing iMGLs after 5-day exposure to 100 vg/cell of BI-RGD1: CAG-mScarlet-NLS with and without Vpx, lentivirus with the same promoter and transgene (CAG-mScarlet-NLS genome, 0.025 pg P24 per cell) and Vpx only. Scale bars = 100 μm. **b,** Microscopy images showing iMGLs after 5-day exposure to 100 vg/cell of BI-RGD1, BI-MPR1 and AAV9 packaging CAG-mScarlet-NLS genome. Scale bars = 100 μm. **c,** Volcano plots showing genes that significantly change expression in iMGLs after exposure to AAV (left panel) and AAV + Vpx (right panel). n=3 replicates. p-values were calculated with Deseq2 using a Wald test and adjusted with Benjamini- Hochberg method. **d,** Gene ontology analysis using Enrichr of genes that change significantly in iMGLs after exposure to AAV. **e,** Flow cytometry quantification of in vitro transduction efficiency of BI-RGD1 containing hIBA1 vs. CAG promoter. Points represent individual wells. Error bars represent SD. Numbers on the x axis represent the amount of virus added in vg/cell. Unpaired, two-tailed Student’s t-test was performed to calculate p- values comparing the two different promoters at different conditions. *** represents p- value<0.001 and * represents p-value<0.05. t-statistics and 95% confidence intervals for the comparisons are as follows: 10K (t (4) = 10.94, 6.64% to 11.16%), 100K (t(4) = 2.243, -5.80% to 54.60%), 1M (t(4) = 3.308, 4.71% to 53.87%), 10K + Vpx (t(4) = 11.59, 12.60% to 20.54%), 100K + Vpx (t(4) = 14.58, 24.18% to 35.56%), 1M + Vpx (t(4) = 9.627, 17.15% to 31.06%). **f,** Microscopy images showing iMGLs after treatment with BI-RGD1: hIBA1- mScarlet-NLS. Scale bars = 100 μm.

**Supplementary Fig. 5:**
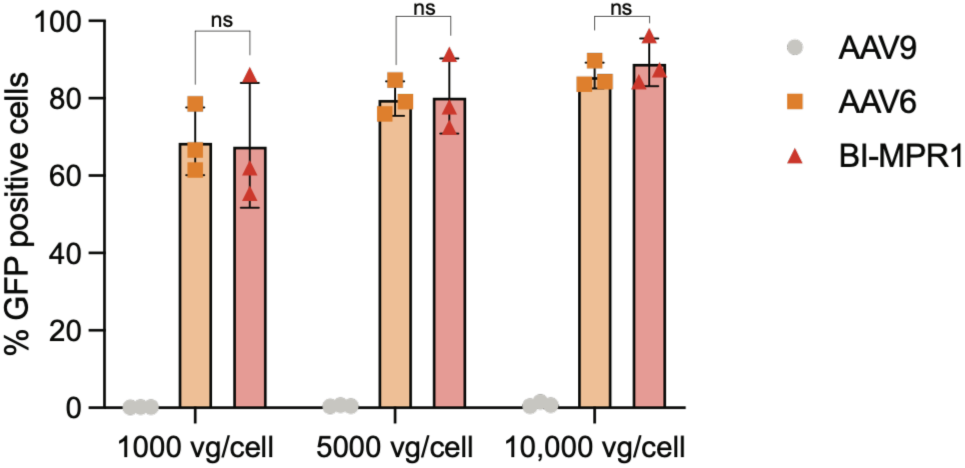
BI-MPR1 with CAG-mScarlet-NLS genome is as efficient at in vitro transduction of primary human T cells as AAV6. Comparison of transduction efficiency in live, CD3+ cells across tested AAV capsids packaging CAG-GFP genome. Points represent individual donors. Error bars represent SD. Statistical significance was assessed using one-way ANOVA with Tukey’s post-hoc multiple-comparisons test. Select post-hoc comparisons are shown.

**Supplementary Fig. 6:**
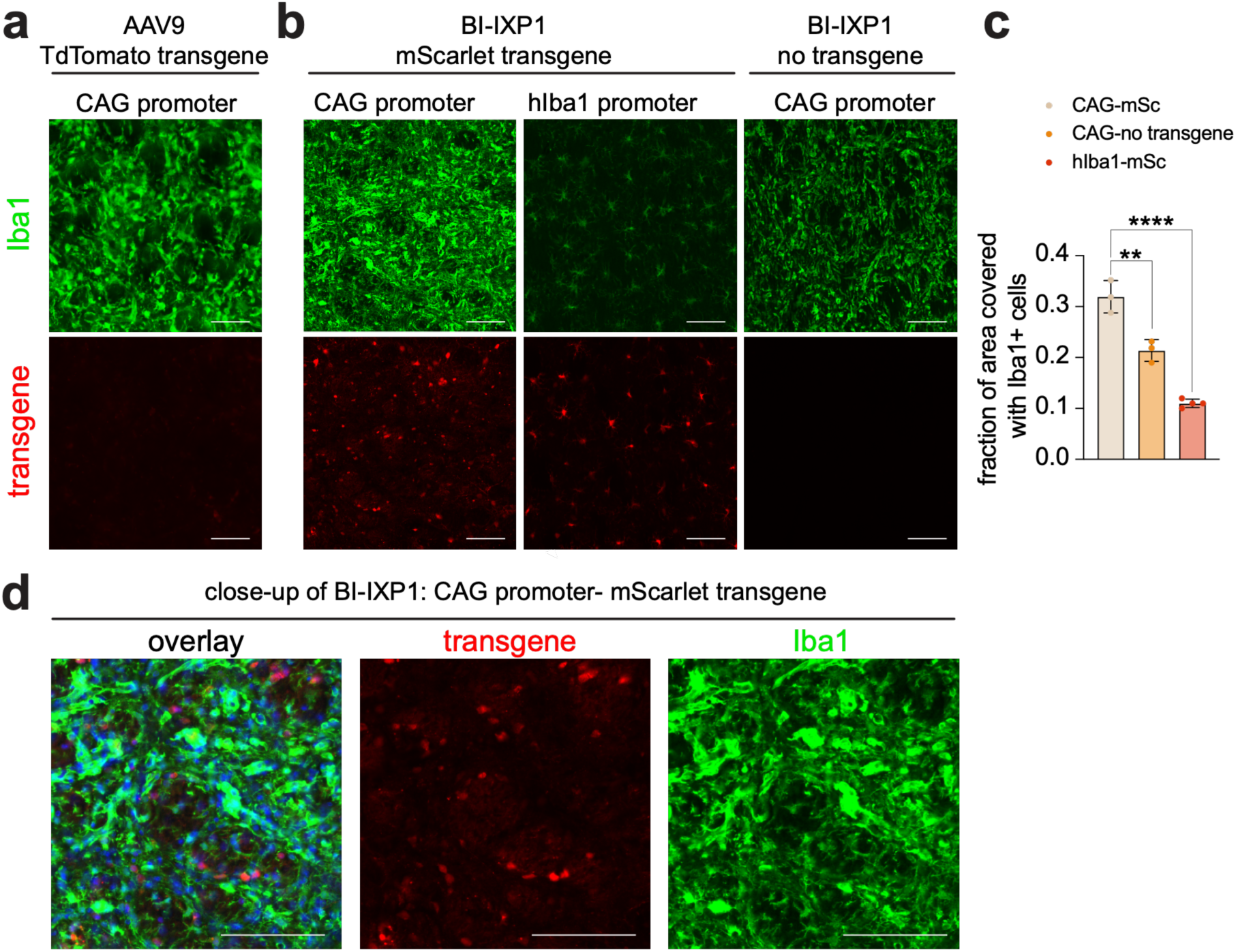
CAG promoter mediates microglia activation in vivo. a, AAV9 with Cre-dependent TdTomato transgene and CAG promoter (CAG-DIO-TdTomato) intrastriatally injected into Cx3cr1-Cre-ER mice (4E9 vg per site). Scale bars = 100 μm. **b,** Direct comparison of AAVs with CAG and hIBA1 promoters expressing mScarlet, packaged in BI-IXP1 capsid. The third panel shows a virus that does not contain a transgene and has pA sites cloned after the CAG promoter to prevent expression from any downstream ORFs **c,** Quantification of area covered with Iba1+ cells shown in B (N=3-4 biological replicates per condition) as a measure of microgliosis. Points represent individual mice. Error bars represent SD. Ordinary one-way ANOVA (F (2, 7) = 84.14, P < 0.0001) with Tukey’s post-hoc multiple-comparisons test was performed to calculate p- values. Select post-hoc comparisons are shown. ** represents p < 0.01 and **** represents p < 0.0001. 95% confidence intervals for the difference between means were 0.16 to 0.26 for CAG-mScarlet vs. hIBA1-mScarlet and 0.06 to 0.15 for CAG-mScarlet vs. CAG-no transgene. **d,** Magnified view of striatum injected with BI-IXP1: CAG-mScarlet- NLS shown in **b.** Scale bars = 100 μm.

**Supplementary Fig. 7:**
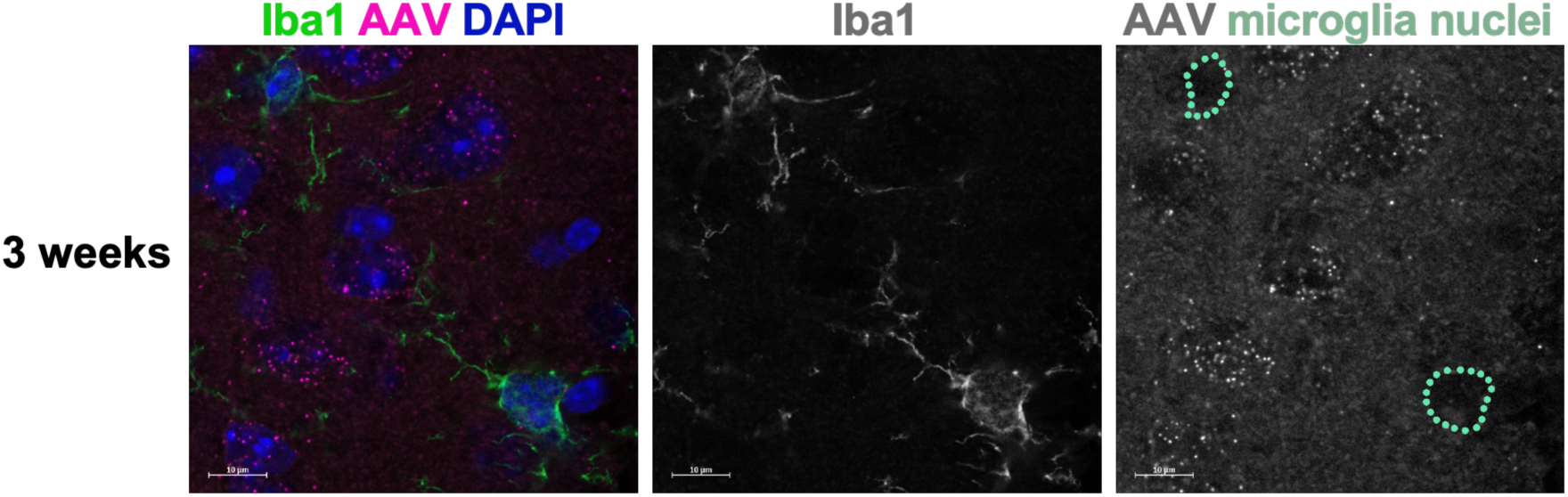
AAV genomes containing hIBA1 promoter and miR-124 target sites preferentially accumulate in non-microglia cells. DNA in situ hybridization for AAV genome DNA was done 3 weeks after intracranial injection of BI-IXP1:hIBA1-mScarlet- miR124TS. Showing 1 optical section. Microglia were stained with Iba1 after DNA in situ hybridization using a probe set specific for WPRE in the viral genome. The panel on the right shows outlines of microglia nuclei in green as defined by DAPI staining. n=3 mice. Scale bars = 10 μm.

**Supplementary Fig. 8:**
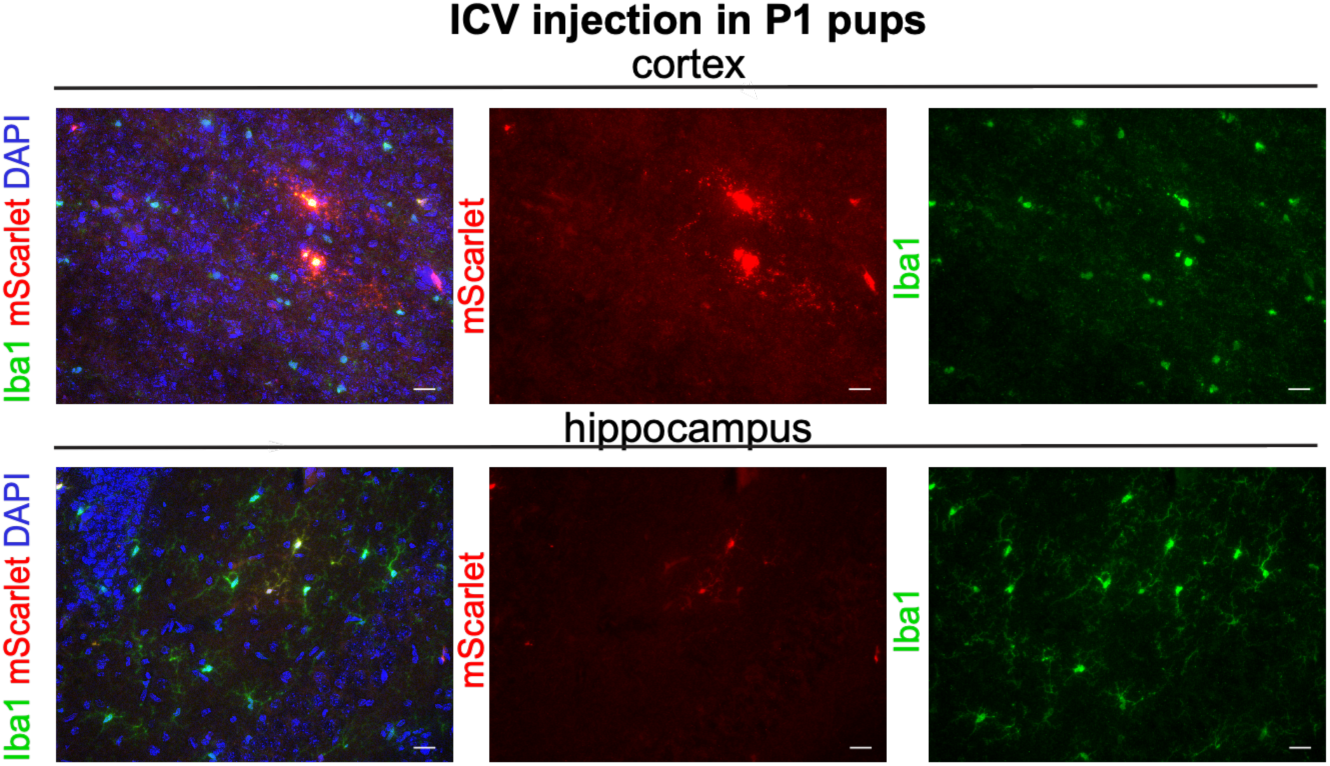
Microglia-targeting AAV injection into cerebral ventricles results in sparse transduction of multiple brain regions. ICV injection of BI-IXP1: hIBA1-mScarlet-miR124TS in P1 pups. 7.78E9 vg per hemisphere. n=3 mice per experiment. Transduction was visualized 3 weeks after injection. mScarlet signal was amplified using an anti-RFP antibody. Scale bars = 50 μm.

## References

1. Abud, E. M., Ramirez, R. N., Martinez, E. S., Healy, L. M., Nguyen, C. H. H., Newman, S. A., Yeromin, A. V., Scarfone, V. M., Marsh, S. E., Fimbres, C., Caraway, C. A., Fote, G. M., Madany, A. M., Agrawal, A., Kayed, R., Gylys, K. H., Cahalan, M. D., Cummings, B. J., Antel, J. P., … Blurton-Jones, M. (2017). iPSC-Derived Human Microglia-like Cells to Study Neurological Diseases. Neuron, 94(2), 278–293.e9.

2. Aoki, R., Konno, A., Hosoi, N., Kawabata, H., & Hirai, H. (2025). AAV vectors for specific and efficient gene expression in microglia. Cell Reports Methods, 5(8), 101116.

3. Bankhead, P., Loughrey, M. B., Fernández, J. A., Dombrowski, Y., McArt, D. G., Dunne, P. D., McQuaid, S., Gray, R. T., Murray, L. J., Coleman, H. G., James, J. A., Salto-Tellez, M., & Hamilton, P. W. (2017). QuPath: Open-source software for digital pathology image analysis. Scientific Reports, 7(1), 16878.

4. Berry, G. E., & Asokan, A. (2016). Cellular transduction mechanisms of adeno-associated viral vectors. Current Opinion in Virology, 21, 54–60.

5. Cao, W., Tan, Z., Berackey, B. T., Nguyen, J. K., Brown, S. R., Du, S., Lin, B., Ye, Q., Seiler, M., Holmes, T. C., & Xu, X. (2025). An AAV capsid proposed as microglia-targeting directs genetic expression in forebrain excitatory neurons. Cell Reports Methods, 5(6), 101054.

6. Chan, K. Y., Jang, M. J., Yoo, B. B., Greenbaum, A., Ravi, N., Wu, W.-L., Sánchez-Guardado, L., Lois, C., Mazmanian, S. K., Deverman, B. E., & Gradinaru, V. (2017). Engineered AAVs for efficient noninvasive gene delivery to the central and peripheral nervous systems. Nature Neuroscience, 20(8), 1172–1179.

7. Crooks, G. E., Hon, G., Chandonia, J.-M., & Brenner, S. E. (2004). WebLogo: a sequence logo generator. Genome Research, 14(6), 1188–1190.

8. Cui, M., Su, Q., Yip, M., McGowan, J., Punzo, C., Gao, G., & Tai, P. W. L. (2024). The AAV2.7m8 capsid packages a higher degree of heterogeneous vector genomes than AAV2. Gene Therapy, 31(9-10), 489–498.

9. Deverman, B. E., Pravdo, P. L., Simpson, B. P., Kumar, S. R., Chan, K. Y., Banerjee, A., Wu, W.-L., Yang, B., Huber, N., Pasca, S. P., & Gradinaru, V. (2016). Cre-dependent selection yields AAV variants for widespread gene transfer to the adult brain. Nature Biotechnology, 34(2), 204–209.

10. Dobin, A., Davis, C. A., Schlesinger, F., Drenkow, J., Zaleski, C., Jha, S., Batut, P., Chaisson, M., & Gingeras, T. R. (2013). STAR: ultrafast universal RNA-seq aligner. *Bioinformatics (Oxford*, England), 29(1), 15–21.

11. Dolan, M.-J., Therrien, M., Jereb, S., Kamath, T., Gazestani, V., Atkeson, T., Marsh, S. E., Goeva, A., Lojek, N. M., Murphy, S., White, C. M., Joung, J., Liu, B., Limone, F., Eggan, K., Hacohen, N., Bernstein, B. E., Glass, C. K., Leinonen, V., … Stevens, B. (2023). Exposure of iPSC-derived human microglia to brain substrates enables the generation and manipulation of diverse transcriptional states in vitro. Nature Immunology, 24(8), 1382– 1390.

12. Drouyer, M., Merjane, J., Nedelkoska, T., Westhaus, A., Scott, S., Lee, S., Burke, P. G. R., McMullan, S., Lanciego, J. L., Vicente, A. F., Bugallo, R., Unzu, C., González-Aseguinolaza, G., Gonzalez-Cordero, A., & Lisowski, L. (2024). Enhanced AAV transduction across preclinical CNS models: A comparative study in human brain organoids with cross-species evaluations. Molecular Therapy. Nucleic Acids, 35(3), 102264.

13. Eyquem, J., Mansilla-Soto, J., Giavridis, T., van der Stegen, S. J. C., Hamieh, M., Cunanan, K. M., Odak, A., Gönen, M., & Sadelain, M. (2017). Targeting a CAR to the TRAC locus with CRISPR/Cas9 enhances tumour rejection. Nature, 543(7643), 113–117.

14. Favuzzi, E., Huang, S., Saldi, G. A., Binan, L., Ibrahim, L. A., Fernández-Otero, M., Cao, Y., Zeine, A., Sefah, A., Zheng, K., Xu, Q., Khlestova, E., Farhi, S. L., Bonneau, R., Datta, S. R., Stevens, B., & Fishell, G. (2021). GABA-receptive microglia selectively sculpt developing inhibitory circuits. Cell, 184(22), 5686.

15. Grimm, D., Lee, J. S., Wang, L., Desai, T., Akache, B., Storm, T. A., & Kay, M. A. (2008). In vitro and in vivo gene therapy vector evolution via multispecies interbreeding and retargeting of adeno-associated viruses. Journal of Virology, 82(12), 5887–5911.

16. Gurtsieva, D., Minskaia, E., Zhuravleva, S., Subcheva, E., Sakhibgaraeva, E., Brovin, A., Tumaev, A., & Karabelsky, A. (2024). Engineered AAV2.7m8 Serotype Shows Significantly Higher Transduction Efficiency of ARPE-19 and HEK293 Cell Lines Compared to AAV5, AAV8 and AAV9 Serotypes. Pharmaceutics, 16(1).

17. Hammond, T. R., Dufort, C., Dissing-Olesen, L., Giera, S., Young, A., Wysoker, A., Walker, A. J., Gergits, F., Segel, M., Nemesh, J., Marsh, S. E., Saunders, A., Macosko, E., Ginhoux, F., Chen, J., Franklin, R. J. M., Piao, X., McCarroll, S. A., & Stevens, B. (2019). Single-Cell RNA Sequencing of Microglia throughout the Mouse Lifespan and in the Injured Brain Reveals Complex Cell-State Changes. Immunity, 50(1), 253–271.e6.

18. Hasselmann, J., Coburn, M. A., England, W., Figueroa Velez, D. X., Kiani Shabestari, S., Tu, C. H., McQuade, A., Kolahdouzan, M., Echeverria, K., Claes, C., Nakayama, T., Azevedo, R., Coufal, N. G., Han, C. Z., Cummings, B. J., Davtyan, H., Glass, C. K., Healy, L. M., Gandhi, S. P., … Blurton-Jones, M. (2019). Development of a Chimeric Model to Study and Manipulate Human Microglia In Vivo. Neuron, 103(6), 1016–1033.e10.

19. Hemmi, H., Takeuchi, O., Kawai, T., Kaisho, T., Sato, S., Sanjo, H., Matsumoto, M., Hoshino, K., Wagner, H., Takeda, K., & Akira, S. (2000). A Toll-like receptor recognizes bacterial DNA. Nature, 408(6813), 740–745.

20. Keren-Shaul, H., Spinrad, A., Weiner, A., Matcovitch-Natan, O., Dvir-Szternfeld, R., Ulland, T. K., David, E., Baruch, K., Lara-Astaiso, D., Toth, B., Itzkovitz, S., Colonna, M., Schwartz, M., & Amit, I. (2017). A Unique Microglia Type Associated with Restricting Development of Alzheimer’s Disease. Cell, 169(7), 1276–1290.e17.

21. Khaparde, A., Patra, R., Roy, S., Babu G R, S., & Ghosh, A. (2025). Improving AAV Production Yield and Quality for Different Serotypes Using Distinct Processing Methods. ACS Omega, 10(22), 22657–22670.

22. Krasemann, S., Madore, C., Cialic, R., Baufeld, C., Calcagno, N., El Fatimy, R., Beckers, L., O’Loughlin, E., Xu, Y., Fanek, Z., Greco, D. J., Smith, S. T., Tweet, G., Humulock, Z., Zrzavy, T., Conde-Sanroman, P., Gacias, M., Weng, Z., Chen, H., … Butovsky, O. (2017). The TREM2-APOE Pathway Drives the Transcriptional Phenotype of Dysfunctional Microglia in Neurodegenerative Diseases. Immunity, 47(3), 566–581.e9.

23. Kunkle, B. W., Grenier-Boley, B., Sims, R., Bis, J. C., Damotte, V., Naj, A. C., Boland, A., Vronskaya, M., van der Lee, S. J., Amlie-Wolf, A., Bellenguez, C., Frizatti, A., Chouraki, V., Martin, E. R., Sleegers, K., Badarinarayan, N., Jakobsdottir, J., Hamilton-Nelson, K. L., Moreno-Grau, S., … Genetic and Environmental Risk in AD/Defining Genetic, Polygenic and Environmental Risk for Alzheimer’s Disease Consortium (GERAD/PERADES),. (2019). Genetic meta-analysis of diagnosed Alzheimer’s disease identifies new risk loci and implicates Aβ, tau, immunity and lipid processing. Nature Genetics, 51(3), 414–430.

24. Liao, Y., Smyth, G. K., & Shi, W. (2014). featureCounts: an efficient general purpose program for assigning sequence reads to genomic features. *Bioinformatics (Oxford*, England), 30(7), 923–930.

25. Ling, Q., Herstine, J. A., Bradbury, A., & Gray, S. J. (2023). AAV-based in vivo gene therapy for neurological disorders. Nature Reviews Drug Discovery, 22(10), 789–806.

26. Lin, R., Zhou, Y., Yan, T., Wang, R., Li, H., Wu, Z., Zhang, X., Zhou, X., Zhao, F., Zhang, L., Li, Y., & Luo, M. (2022). Directed evolution of adeno-associated virus for efficient gene delivery to microglia. Nature Methods, 19(8), 976–985.

27. Li, Q., Cheng, Z., Zhou, L., Darmanis, S., Neff, N. F., Okamoto, J., Gulati, G., Bennett, M. L., Sun, L. O., Clarke, L. E., Marschallinger, J., Yu, G., Quake, S. R., Wyss-Coray, T., & Barres, B. A. (2019). Developmental Heterogeneity of Microglia and Brain Myeloid Cells Revealed by Deep Single-Cell RNA Sequencing. Neuron, 101(2), 207–223.e10.

28. Love, M. I., Huber, W., & Anders, S. (2014). Moderated estimation of fold change and dispersion for RNA-seq data with DESeq2. Genome Biology, 15(12), 550.

29. Luo, N., Lin, K., Cai, Y., Sui, X., Zhang, Z., Xing, J., Liu, G., Yuan, W., Wang, J., & Xu, F. (2024). Microglia-specific transduction via AAV11 armed with IBA1 promoter and miRNA-9 targeting sequences. *bioRxiv*, July 9.

30. Maes, M. E., Colombo, G., Schulz, R., & Siegert, S. (2019). Targeting microglia with lentivirus and AAV: Recent advances and remaining challenges. Neuroscience Letters, 707, 134310.

31. Morshed, N., Demers, M., Gonzalez-Ramos, A., Jäntti, H., Doman, J., D’Souza, S., Li, L., Granger, A. J., Johnson, M. B., & Stevens, B. (2026). iAstrocytes model cytokine influences on complement expression and neuronal network synchronization. bioRxiv, June 4.

32. Nayak, S., & Herzog, R. W. (2010). Progress and prospects: immune responses to viral vectors. Gene Therapy, 17(3), 295–304.

33. Nehme, R., Zuccaro, E., Ghosh, S. D., Li, C., Sherwood, J. L., Pietilainen, O., Barrett, L. E., Limone, F., Worringer, K. A., Kommineni, S., Zang, Y., Cacchiarelli, D., Meissner, A., Adolfsson, R., Haggarty, S., Madison, J., Muller, M., Arlotta, P., Fu, Z., Feng, G., & Eggan, K. (2018). Combining NGN2 programming with developmental patterning generates human excitatory neurons with NMDAR-mediated synaptic transmission. Cell Reports, 23(8), 2509–2523.

34. Nonnenmacher, M., Wang, W., Child, M. A., Ren, X.-Q., Huang, C., Ren, A. Z., Tocci, J., Chen, Q., Bittner, K., Tyson, K., Pande, N., Chung, C. H.-Y., Paul, S. M., & Hou, J. (2021). Rapid evolution of blood-brain-barrier-penetrating AAV capsids by RNA-driven biopanning. Molecular Therapy. Methods & Clinical Development, 20, 366–378.

35. Nonnenmacher, M., & Weber, T. (2011). Adeno-associated virus 2 infection requires endocytosis through the CLIC/GEEC pathway. Cell Host & Microbe, 10(6), 563–576.

36. Okada, Y., Hosoi, N., Matsuzaki, Y., Fukai, Y., Hiraga, A., Nakai, J., Nitta, K., Shinohara, Y., Konno, A., & Hirai, H. (2022). Development of microglia-targeting adeno-associated viral vectors as tools to study microglial behavior in vivo. Communications Biology, 5(1), 1224.

37. Paloneva, J., Manninen, T., Christman, G., Hovanes, K., Mandelin, J., Adolfsson, R., Bianchin, M., Bird, T., Miranda, R., Salmaggi, A., Tranebjaerg, L., Konttinen, Y., & Peltonen, L. (2002). Mutations in two genes encoding different subunits of a receptor signaling complex result in an identical disease phenotype. American Journal of Human Genetics, 71(3), 656– 662.

38. Parkhurst, C. N., Yang, G., Ninan, I., Savas, J. N., Yates, J. R., 3rd, Lafaille, J. J., Hempstead, B. L., Littman, D. R., & Gan, W.-B. (2013). Microglia promote learning-dependent synapse formation through brain-derived neurotrophic factor. Cell, 155(7), 1596–1609.

39. Pillay, S., Meyer, N. L., Puschnik, A. S., Davulcu, O., Diep, J., Ishikawa, Y., Jae, L. T., Wosen, J. E., Nagamine, C. M., Chapman, M. S., & Carette, J. E. (2016). An essential receptor for adeno-associated virus infection. Nature, 530(7588), 108–112.

40. Rademakers, R., Baker, M., Nicholson, A. M., Rutherford, N. J., Finch, N., Soto-Ortolaza, A., Lash, J., Wider, C., Wojtas, A., DeJesus-Hernandez, M., Adamson, J., Kouri, N., Sundal, C., Shuster, E. A., Aasly, J., MacKenzie, J., Roeber, S., Kretzschmar, H. A., Boeve, B. F., … Wszolek, Z. K. (2011). Mutations in the colony stimulating factor 1 receptor (CSF1R) gene cause hereditary diffuse leukoencephalopathy with spheroids. Nature Genetics, 44(2), 200–205.

41. Rice, G. I., Bond, J., Asipu, A., Brunette, R. L., Manfield, I. W., Carr, I. M., Fuller, J. C., Jackson, R. M., Lamb, T., Briggs, T. A., Ali, M., Gornall, H., Couthard, L. R., Aeby, A., Attard- Montalto, S. P., Bertini, E., Bodemer, C., Brockmann, K., Brueton, L. A., … Crow, Y. J. (2009). Mutations involved in Aicardi-Goutières syndrome implicate SAMHD1 as regulator of the innate immune response. Nature Genetics, 41(7), 829–832.

42. Safaiyan, S., Besson-Girard, S., Kaya, T., Cantuti-Castelvetri, L., Liu, L., Ji, H., Schifferer, M., Gouna, G., Usifo, F., Kannaiyan, N., Fitzner, D., Xiang, X., Rossner, M. J., Brendel, M., Gokce, O., & Simons, M. (2021). White matter aging drives microglial diversity. Neuron, 109(7), 1100–1117.e10.

43. Santoscoy, M. C., Espinoza, P., Hanlon, K. S., Yang, L., Nieland, L., Ng, C., Badr, C. E., Hickman, S., de la Cruz, D., Griciuc, A., Elkhoury, J., Bennett, R. E., Shen, S., & Maguire, C. A. (2024). Expression-based selection identifies a microglia-tropic AAV capsid for direct and CSF routes of administration in mice. bioRxiv, September 24.

44. Schafer, D. P., Lehrman, E. K., Kautzman, A. G., Koyama, R., Mardinly, A. R., Yamasaki, R., Ransohoff, R. M., Greenberg, M. E., Barres, B. A., & Stevens, B. (2012). Microglia sculpt postnatal neural circuits in an activity and complement-dependent manner. Neuron, 74(4), 691–705.

45. Serrano, C., Cananzi, S., Shen, T., Wang, L.-L., & Zhang, C.-L. (2024). Simple and highly specific targeting of resident microglia with adeno-associated virus. iScience, 27(9), 110706.

46. Stirling, D. R., Swain-Bowden, M. J., Lucas, A. M., Carpenter, A. E., Cimini, B. A., & Goodman, A. (2021). CellProfiler 4: improvements in speed, utility and usability. BMC Bioinformatics, 22(1), 433.

47. Tabebordbar, M., Lagerborg, K. A., Stanton, A., King, E. M., Ye, S., Tellez, L., Krunnfusz, A., Tavakoli, S., Widrick, J. J., Messemer, K. A., Troiano, E. C., Moghadaszadeh, B., Peacker, B. L., Leacock, K. A., Horwitz, N., Beggs, A. H., Wagers, A. J., & Sabeti, P. C. (2021). Directed evolution of a family of AAV capsid variants enabling potent muscle-directed gene delivery across species. Cell, 184(19), 4919–4938.e22.

48. Velasco, S., Kedaigle, A. J., Simmons, S. K., Nash, A., Rocha, M., Quadrato, G., Paulsen, B., Nguyen, L., Adiconis, X., Regev, A., Levin, J. Z., & Arlotta, P. (2019). Individual brain organoids reproducibly form cell diversity of the human cerebral cortex. Nature, 570(7762), 523–527.

49. Wang, J., DeClercq, J. J., Hayward, S. B., Li, P. W.-L., Shivak, D. A., Gregory, P. D., Lee, G., & Holmes, M. C. (2016). Highly efficient homology-driven genome editing in human T cells by combining zinc-finger nuclease mRNA and AAV6 donor delivery. Nucleic Acids Research, 44(3), e30.

50. Weinmann, J., Weis, S., Sippel, J., Tulalamba, W., Remes, A., El Andari, J., Herrmann, A.-K., Pham, Q. H., Borowski, C., Hille, S., Schönberger, T., Frey, N., Lenter, M., VandenDriessche, T., Müller, O. J., Chuah, M. K., Lamla, T., & Grimm, D. (2020). Identification of a myotropic AAV by massively parallel in vivo evaluation of barcoded capsid variants. Nature Communications, 11(1), 5432.

51. Xie, Z., Bailey, A., Kuleshov, M. V., Clarke, D. J. B., Evangelista, J. E., Jenkins, S. L., Lachmann, A., Wojciechowicz, M. L., Kropiwnicki, E., Jagodnik, K. M., Jeon, M., & Ma’ayan, A. (2021). Gene Set Knowledge Discovery with Enrichr. Current Protocols, 1(3), e90.

52. Zhao, J., Yue, Y., Patel, A., Wasala, L., Karp, J. F., Zhang, K., Duan, D., & Lai, Y. (2020). High-Resolution Histological Landscape of AAV DNA Distribution in Cellular Compartments and Tissues following Local and Systemic Injection. Molecular Therapy - Methods & Clinical Development, 18, 856–868.

53. Zhu, J., Huang, X., & Yang, Y. (2009). The TLR9-MyD88 pathway is critical for adaptive immune responses to adeno-associated virus gene therapy vectors in mice. The Journal of Clinical Investigation, 119(8), 2388–2398.

